# Loop 1 of APOBEC3C regulates its antiviral activity against HIV-1

**DOI:** 10.1101/2020.02.05.936021

**Authors:** Ananda Ayyappan Jaguva Vasudevan, Kannan Balakrishnan, Christoph G. W. Gertzen, Fanni Borvető, Zeli Zhang, Anucha Sangwiman, Ulrike Held, Caroline Küstermann, Sharmistha Banerjee, Gerald G. Schumann, Dieter Häussinger, Ignacio G. Bravo, Holger Gohlke, Carsten Münk

## Abstract

APOBEC3 deaminases (A3s) provide mammals with an anti-retroviral barrier by catalyzing dC-to-dU deamination on viral ssDNA. Within primates, A3s have evolved diversely *via* gene duplications and fusions. Human APOBEC3C (hA3C) efficiently restricts the replication of viral infectivity factor (*vif*)-deficient *Simian immunodeficiency virus* (SIVΔ*vif*), but for unknown reasons, it inhibits HIV-1Δ*vif* weakly. In catarrhines (Old World monkeys and apes), the A3C loop 1 displays the conserved amino acid pair WE, while the corresponding consensus sequence in A3F and A3D is the largely divergent pair RK, which is also the inferred ancestral sequence for the last common ancestor of A3C|D|F in primates. Here, we report that modifying the WE residues in hA3C loop 1 to RK leads to stronger interactions with ssDNA substrate, facilitating catalytic function, which resulted in a drastic increase in both deamination activity and the ability to restrict HIV-1 and LINE-1 replication. Conversely, the modification hA3F_WE resulted only in a marginal decrease in HIV-1Δ*vif* inhibition. The two series of ancestral gene duplications that generated A3C, A3D-CTD and A3F-CTD allowed neo/subfunctionalization: A3F-CTD maintained the ancestral RK residues in loop 1, while strong evolutionary pressure selected for the RK→WE modification in catarrhines A3C, possibly allowing for novel substrate specificity and function.

**AUTHOR SUMMARY:** The restriction factors of the APOBEC3 (A3) family of cytidine deaminases inhibit the replication of Vif-deficient retroviruses mainly by mutating their viral genomes. While there are seven A3 proteins (A3A-A3H) found in humans only A3G and A3F potently inhibit HIV-1 replication. A3C in general and its retroviral restriction capacity have not been widely studied probably due to its weak anti-HIV-1 activity, however, it displays a strong antiviral effect against SIV. Understanding the role of A3C is important because it is highly expressed in CD4+ T cells, is upregulated upon HIV-1 infection, and is distributed cell-wide. In this study, we report that replacing two residues in loop 1 of A3C protein with conserved positively-charged amino acids enhance the substrate DNA binding, which markedly facilitates its deamination-dependent antiviral activity against HIV-1 as well as increasing the restriction of LINE-1 retroelements. Furthermore, our evolutionary analysis demonstrates that the pressure that caused the loss of potential loop 1 residues occurred only in A3C but not in primate homologues. Overall, our study highlights the possibility of A3C acting as a super restriction factor, however, this was likely evolutionarily selected against to achieve a balance between anti-viral/anti-LINE-1 activity and genotoxicity.

## INTRODUCTION

The APOBEC3 (A3) family of single-stranded (ss) DNA cytidine deaminases builds an intrinsic immune defense against retroviruses, retrotransposons, and other viral pathogens [1–4]. There are seven human A3 proteins (A3s) that possess either one (A3A, A3C, and A3H) or two (A3B, A3D, A3F, and A3G) zinc (Z)-coordinating DNA cytosine deaminase motifs, HXE[X_23-28_]PC[X_2-4_]C (where X indicates a non-conserved position) [5–7]. A3G was identified as a factor capable of restricting infection of HIV-1 lacking Vif (viral infectivity factor) protein in non-permissive T cell lines whose biochemical properties and biological functions were extensively studied [3,8–11].

The encapsidation of A3 into the viral particles is crucial for virus inhibition [12–17]. During reverse transcription, viral core-associated A3 enzymes can deaminate cytidines (dC) on the retroviral ssDNA into uridines (dU). These base modifications in the minus DNA strand cause coding changes and premature stop codons in the plus-strand viral genome (dG→dA hypermutation), which impair or suppress viral infectivity [2,9,18–21]. In addition to the mutagenic activity of the viral-incorporated A3 enzyme, deaminase-independent mechanisms of restriction were also manifested by impeding reverse transcription or inhibiting DNA integration [22–27]. To counteract A3 mediated inhibition, lentiviruses evolved the Vif protein, which physically interacts with A3s to target them for polyubiquitination and proteasomal degradation, and thereby depleting the cellular A3s [28–30]. These A3-Vif interactions are often species-specific [31–35].

A3D, A3F, A3G, and A3H were shown to restrict HIV-1 lacking *vif* (HIV-1Δ*vif*) [2,35–39]. Recently, mutation signatures resulting from the catalytic activity of nuclear localized A3s (especially A3A, A3B, and likely A3H) were reported in several cancer types [40, 41] (for reviews, see: [42–45]. However, the A3C, which is distributed in both cytoplasm and nucleus [46] seems not to be a causative agent of chromosomal DNA mutations. Human A3C is known to act as a potent inhibitor of *Simian immunodeficiency virus* from African green monkey (SIVagm) and SIVmac (from rhesus macaque), limits the infectivity of herpes simplex virus, certain human papillomaviruses, murine leukemia virus, Bet-deficient foamy virus, and hepatitis B virus and represses the replication of LINE-1 (L1) retrotransposons [46–56]. However, the restrictive role of A3C on HIV-1 is marginal and there are several contradictory findings regarding its viral packaging and cytidine deamination activity [39,47,57–59]. Notably, A3C is expressed ubiquitously in lymphoid cells [5,47,60,61]. mRNA expression levels of A3C were found to be higher in HIV-infected CD4^+^ T lymphocytes [39, 47], and significantly elevated in elite controllers with respect to ART-suppressed individuals [62]. A3C was found to moderately deaminate HIV-1 DNA if expressed in target cells of the virus and rather increased viral diversity than caused restriction [60].

The crystal structure of A3C and its HIV-1 Vif-binding interface were reported recently [63]. The study revealed several key residues in the hydrophobic V-shaped groove formed by the α2 and α3 helices of A3C that facilitate Vif binding resulting in proteasome-mediated degradation of A3C [63]. We extended this finding and identified additional Vif interaction sites in α4 helix of A3C [64]. Other than a previous study that predicted putative DNA substrate binding pockets [52], biochemical and structural aspects of A3C enzymatic activity and their relevance for antiviral activity are not well investigated to date [3, 4].

Recently, we have shown that increasing the catalytic activity of A3C by an S61P substitution (based on the structural homology found between A3C and A3F at their C-terminal domain, A3F-CTD) is not sufficient to inhibit HIV-1Δ*vif* [65]. It is unclear why A3C can potently restrict SIVΔ*vif*, but not HIV-1Δ*vif* despite the fact that the wild-type human enzyme possesses reasonable catalytic activity and encapsidates efficiently into retroviral particles [65]. Here we set out to understand the function of A3C in the context of HIV-1 inhibition. We generated a synthetic open reading frame derived from sooty mangabey monkey genome *(smm, Cercocebus atys (torquatus) lunulatus),* encoding for an A3C-like protein (hereafter called smmA3C-like protein) capable of restricting HIV-1 to similar or higher extent than human A3G. This A3C-like protein was reported to be resistant to HIV-1 Vif-mediated depletion [64]. Using this smmA3C-like protein as a tool, here we dissect the structure-function of hA3C and identify the crucial regions of A3C that facilitate stronger inhibition of HIV-1.

## RESULTS

### Identification of an A3Z2 protein with enhanced antiviral activity

To determine whether A3C from non-human primates can potently restrict HIV-1Δ*vif* propagation, we produced HIV-1Δ*vif* luciferase reporter virus particles with A3C (an A3Z2 protein) from human, rhesus macaque, chimpanzee (cpz), African green monkey (agm), with human A3G (an A3Z2-Z1, double domain protein), or a synthetic smmA3C-like protein and tested their viral infectivity. Viral particles were pseudotyped with the glycoprotein of *Vesicular stomatitis virus* (VSV-G) and normalized by reverse transcriptase (RT) activity before infection. The luciferase enzyme activity of infected cells was quantified two days post infection. Figure 1A shows the level of relative infectivity of HIV-1Δ*vif* in the presence of the tested A3C proteins and hA3G. Human, rhesus, chimpanzee, and African green monkey A3C proteins reduced the relative infectivity of HIV-1Δ*vif* similarly by approximately 60 to 70%.

**Figure 1.**
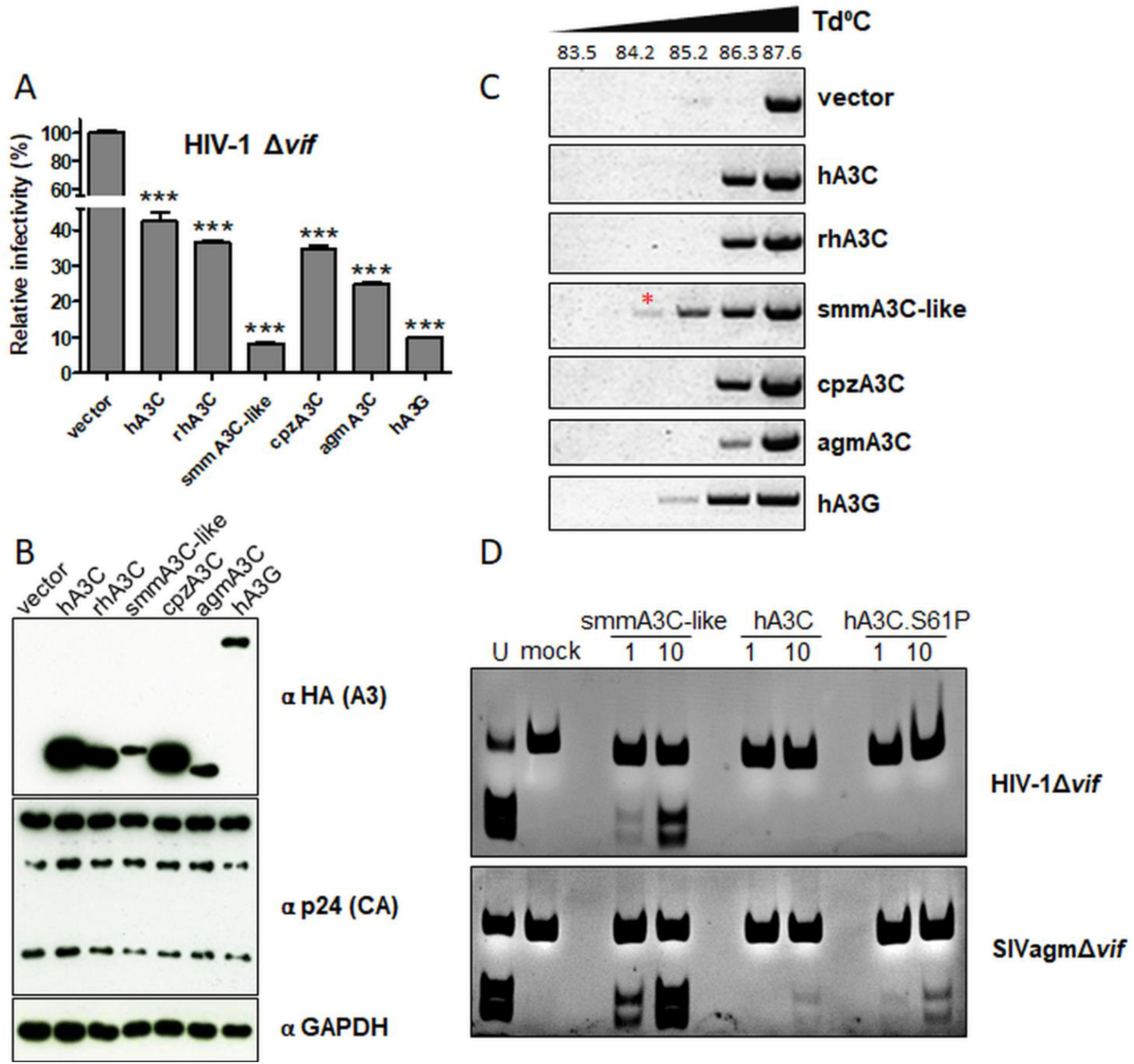
A3C-like protein from sooty mangabey but not A3C from any orthologue inhibits HIV1Δ*vif* by more than an order of magnitude. (A) HIV-1Δ*vif* particles were produced with A3C from human, rhesus macaque, chimpanzees (cpz), African green monkey (agm), and A3C-like protein from sooty mangabey monkey (smm), hA3G or vector only. Infectivity of (RT-activity normalized) equal amounts of viruses, relative to the virus lacking any A3, was determined by quantification of luciferase activity in HEK293T cells. Presented values are means ± standard deviations (error bars) for three independent experiments. Unpaired t-tests were computed to determine whether differences between vector and each A3 protein reach the level of statistical significance. Asterisks indicate statistically significant differences: ***, *p* < 0.0001. (B) Immunoblot analysis of HA-tagged A3 and HIV-1 capsid expression in cell lysates using anti-HA and anti p24 antibodies, respectively. GAPDH served as a loading control. “α” represents anti. Viral encapsidation of hA3G, hA3C, and smmA3C-like protein is demonstrated in Suppl. Fig. S1. (C) 3D-PCR: HIV-1Δ*vif* produced together with A3C orthologues, hA3G or vector controls were used to transduce HEK293T cells. Total DNA was extracted and a 714-bp fragment of reporter viral DNA was selectively amplified using 3D-PCR. T_d_ = denaturation temperature. Extensive viral DNA editing profile of smmA3C-like protein and its relative positions of G→A transition mutations are presented in Suppl. Fig. S5. (D) *In vitro* deamination activity of A3Cs encapsidated in HIV-1Δ*vif*, and SIVagmΔ*vif* particles. Virions were concentrated and lysed in mild lysis buffer and an equal amount of lysate were used for the assay. Numbers 1 and 10 indicate 60 ng and 600 ng of A3 expression vector used for transfection, respectively. Samples were treated with RNAse A; oligonucleotide-containing uracil (U) instead of cytosine served as a marker to denote the migration of deaminated product after restriction enzyme cleavage. S-substrate, P-product.

Conversely, smmA3C-like protein inhibited HIV-1Δ*vif* replication by more than one order of magnitude (Fig. 1A). Human A3G served as a positive control. Expression of the A3s in viral vector-producing cells showed that expression levels of smmA3C-like protein and agmA3C were lower than those of A3Cs from human, rhesus, and cpz (Fig. 1B). Viral incorporation of smmA3C-like protein was found to be very similar to hA3G, but much less efficient compared to hA3C (Suppl. Fig. S1A).

The smmA3C-like construct was originally described to express A3C of smm [64]. However, using alignments of primate A3Z2 and related A3 proteins, we later found that the generated open reading frame consists of exons encoded by genes of smmA3C and smmA3F. In the smmA3C-like construct, the first “exon” (encoding for amino acids ^1^MNPQIR^6^) and last “exon” (encoding for amino acids ^153^FKYC to EILE^190^) were derived from smmA3C (smmA3C exon 1 and exon 4) while the second “exon” (encoding for amino acids ^7^NPMK to FRNQ^58^) and third “exon” (encoding for amino acids ^59^VDPE to VDPE^151^) in smmA3C-like were of smmA3F origin (smmA3F C-terminal domain, CTD, exon 5 and exon 6) (Suppl. Fig. S2). Poor annotation of the smm genome and the high sequence similarity let us fuse these exons that were derived from smmA3C and smmA3F during the amplification step. To compare smmA3C-like to the wild-type proteins, we cloned the genuine smmA3C and smmA3F-CTD and tested their activity. We found that only the smmA3C-like protein and not smmA3C protein showed enhanced cytidine deaminase activity (Suppl. Fig. S3). The smmA3F-CTD construct failed to express detectable levels of protein in transfected cells (Suppl. Fig. S3).

To study G-to-A mutations on the plus-strand of viral DNA triggered by A3 *in vivo* activity, we routinely used a method called “3D-PCR” [65, 66]. DNA sequences in which the cytosines were deaminated by A3 activity contain less GC base pairs than non-edited DNA, resulting in a lower melting temperature than the original, non-edited DNA. Therefore, successful amplification at lower denaturation temperatures (T_d_) (83.5 - 87.6°C) by 3D-PCR is indicating the presence of A3-edited sequences. Because restriction of HIV-1Δ*vif* by smmA3C-like protein was similar or slightly stronger than restriction by hA3G (Fig. 1A), we analyzed the DNA editing capacity of these A3s during infection by 3D-PCR on the viral genome. 3D-PCR amplification with samples of cells infected with HIV-1Δ*vif* viruses encapsidating hA3C, rhA3C, cpzA3C, or agmA3C yielded amplicons until T_d_ 86.3°C, whereas the activity of smmA3C-like protein on the same substrate allowed to produce amplicons at lower T_d_, 84.2°C. In control reactions using virions produced in the presence of hA3G, PCR amplification of viral DNA was detectable at lower T_d_ (85.2°C and weakly at 84.2°C) (Fig. 1C). Importantly, using the vector control sample (no A3), PCR amplicons could be amplified only at higher T_d_ (87.6°C). In our previous study, we elaborately compared the hypermutation load and patterns induced by A3C, A3G, and A3F in retroviruses and found that in HIV-1Δ*vif,* the G→A mutation rate induced by hA3C and A3C.S61P (Suppl. Fig. S4, an A3C mutant with enhanced deamination activity against SIVagmΔ*vif*) was about 6%, whereas A3G and A3F triggered mutation rate was above 15% [65]. To study the effect of smmA3C-like protein in HIV-1Δ*vif*, PCR products generated on smmA3C-like protein-edited samples formed at 84.2°C were cloned and independent clones were sequenced. smmA3C-like protein caused hypermutation in HIV-1Δ*vif* with a rate of 17.16% and predominantly favored the expected <u>G</u>A dinucleotide context (Suppl. Fig. S5). In addition, we have applied qualitative *in vitro* cytidine deamination assays using A3 proteins isolated from HIV-1Δ*vif* and SIVagmΔ*vif* viral particles [67, 68]. This PCR-based assay depends on the sequence change caused by A3 converting a dC→dU in an 80-nucleotide (nt) ssDNA substrate harboring the A3C-specific TTCA motif. Catalytic deamination of dC→dU by A3C is then followed by a PCR that replaces dU by dT generating an MseI restriction site. The efficiency of MseI digestion was monitored by using a similar 80-nt substrate containing dU instead of dC in the recognition site. As expected, hA3C and hA3C.S61P, encapsidated into the HIV-1Δ*vif* particles, did not yield a considerable product resulting from ssDNA cytidine deamination [65], however, smmA3C-like protein formed high amounts of deamination products (Fig. 1D). Using smmA3C-like protein, the deamination products were observed even after transfection of 10-fold smaller amounts of expression plasmid during virus production. In contrast, A3C and A3C.S61P proteins isolated from SIVagmΔ*vif* particles but not from HIV-1Δ*vif* particles produced the expected deamination products, whereas smmA3C-like protein exhibited the strongest catalytic activity, regardless of the source (Fig. 1D). Taken together, we conclude that smmA3C-like protein inhibits HIV-1 by cytidine deamination causing hypermutation of the viral DNA.

### Identification of the regulatory domain of smmA3C-like protein that mediates HIV-1 restriction

Amino acid sequence identity and similarity between hA3C and smmA3C-like protein reach 77.9% and 90%, respectively (Suppl. Fig. S4A). To facilitate the identification of distinct determinants of smmA3C-like protein that confer HIV-1 inhibition, ten different hA3C/smmA3C-like chimeras were constructed [64] (Fig. 2A). We first tested the anti-HIV-1Δ*vif* activity of these A3C chimeras. Viral particles containing different chimeric proteins were produced and their infectivity was tested. As shown in Fig. 2B, chimeras C2, C4, and C8 strongly reduced the infectivity of HIV-1Δ*vif*. Especially, chimera C2 (hA3C harboring a swap of 36 residues of the smmA3C-like protein at the N-terminal end) inhibited HIV-1Δ*vif* replication by about two orders of magnitude. On the contrary, chimeras C6 and C9 reduced viral infectivity by 72% relative to vector control (Fig. 2B).

**Figure 2.**
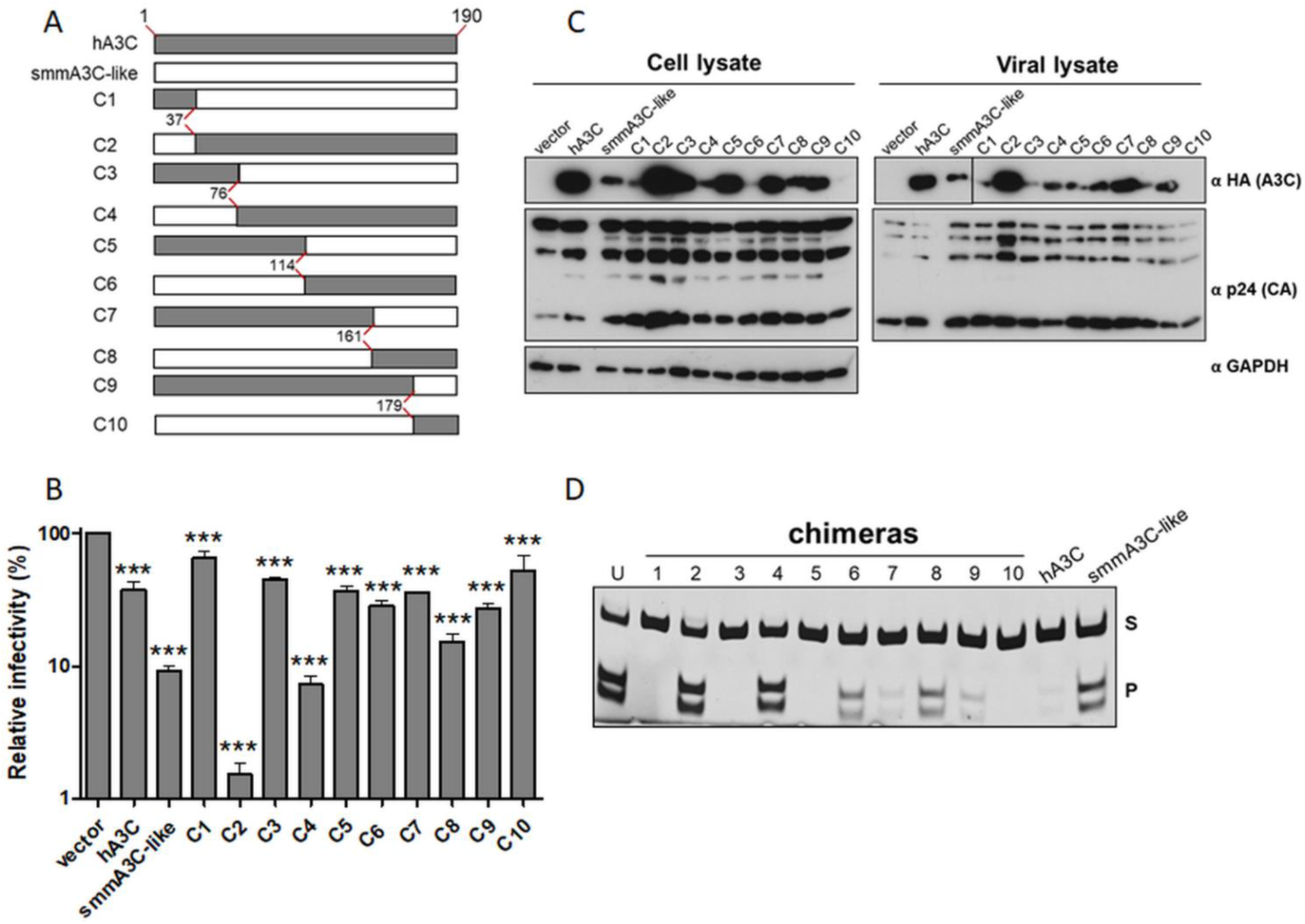
Anti-HIV-1 activity of hA3C/smmA3C-like protein chimeras. (A) Structures of the generated chimeras between A3C and smmA3C-like protein. Grey and white boxes indicate fractions of A3C and the smmA3C-like protein, respectively. Each chimera (C) encompasses 190 amino acids. Amino acid position (number) at the breakpoints of each chimera is indicated. Please find protein sequence and alignment presented in Suppl. Fig. S4A. (B) HIV-1Δ*vif* particles were produced with A3C from human, smm (A3C-like), and h/smm chimeras or vector only. Infectivity of (RT-activity normalized) equal amounts of viruses, relative to the virus lacking any A3, was determined by quantification of luciferase activity in HEK293T cells. Values are means ± standard deviations (error bars) for three independent experiments. Unpaired t-tests were computed to determine whether differences between vector and each A3 protein reach the level of statistical significance. Asterisks represent statistically significant differences: ***, *p* < 0.0001. (C) A3 expression in the cell lysates of transfected cells and A3 viral incorporation were determined by immunoblotting. A3s and HIV-1 capsids were stained with anti-HA and anti-p24 antibodies, respectively. GAPDH served as a loading control. “α” represents anti. (D) To examine the catalytic activity of A3C chimeras, *in vitro* deamination assays were performed using lysates of cells that were previously transfected with respective expression plasmids. RNAse-treatment was included in samples (ten h/smm A3C chimeras, hA3C and smmA3C-like protein) subjected to this assay; oligonucleotide containing uracil (U) instead of cytosine served as a marker to denote the migration of deaminated product after restriction enzyme cleavage. S-substrate, P-product.

Next, we determined the intracellular expression and virion incorporation efficiency of the chimeras by immunoblotting. Chimeras C2, C3, C5, C7, and C9, which contain residues 37 to 76 of hA3C (Fig. 2A), were more highly expressed than C1, C4, C6, and C10 (Fig. 2C). Specifically, chimera C2 displayed higher protein levels than hA3C while C10 was below the detection threshold. Chimeras, C2, C4, C6, C7, and C9 were found to be encapsidated in HIV-1Δ*vif* (Fig. 2C, viral lysate). In particular, C3 and C5 were less efficiently packaged into viral particles although they were present at higher intracellular expression levels. Conversely, C6 produced less protein but its viral incorporation was higher than that of C3 or C5. In addition, we analyzed the *in vitro* cytidine deaminase activity of these chimeras as described above (Fig. 2D). Here we used lysates of transfected HEK293T cells to readily evaluate the catalytic activity of the chimeric A3Cs. As demonstrated in Fig. 2D, only the amounts of deamination products predominantly generated by C2 and C4 were similar to those produced by smmA3C-like protein.

Taken together, chimeras C2 and C4 strongly restricted HIV-1Δ*vif* and are characterized by corresponding *in vitro* deamination activity. C6, by contrast, lost any antiviral and deamination activity, suggesting that the N-terminal region of smmA3C-like protein is crucially involved in the antiviral mechanism. We speculate that residues in C2 and C4 that are absent in C6 complemented the restriction activity of these chimeras. Due to its superior antiviral activity we mainly focused on chimera C2 in our following experiments.

### Synergistic effects of residues in the RKYG motif of chimera C2 and smmA3C-like protein govern their potent antiviral activity

To identify the specific residues in C2 that are essential for its anti-HIV-1 activity, we targeted two N-terminal motifs of C2, namely ^13^DPHIFYFH^20^ (shortly “DHIH”) and ^24^LRKAYG^29^ (named “RKYG”) as presented in the sequence alignments of Suppl. Fig. S4A, and generated more variants of C2 by swapping one, two, or four amino acids with the analogous residues of hA3C as presented in Fig. 3A. First, we cloned the C2 variants C2.DH-YG (YGTQ motif of helix α1) and C2.RKYG-WEND (WEND motif of loop 1, see A3C alignment and ribbon diagram Suppl. Fig. S4) and tested their anti-HIV-1 and deamination activity. This pilot experiment revealed that loop 1 motif RKYG but not α1 helix motif DHIH in C2 is essential for its activity (data not shown). Hence, we constructed the mutants C2.R25W, K26E, Y28N, and G29D (Fig. 3A) and tested them for catalytic and antiviral activity. Since the *in vitro* deaminase activity of the chimeras C1 to C10 correlated with their antiviral activity (Fig. 2B and 2D), we expressed these variants of C2 in HEK293T cells and performed *in vitro* deamination assays. The results of the deamination assay clearly demonstrated that the DH motif in C2 is not relevant for its potent catalytic activity as the C2.DH-YG acted similar to C2 (Fig. 3B), but mutation of the RKYG motif in the RKYG-WEND variant resulted in a loss of deamination activity (Fig. 3B). Interestingly, none of the single amino acid changes in RKYG (C2.R25W, K26E, Y28N, and G29D) resulted in the loss-of-function of C2, albeit the catalytic activities of R25W and K26E were partially reduced (Fig. 3B). Consistent with the data obtained from the *in vitro* assay, the chimeric C2.RKYG-WEND variant failed to restrict the infectivity of HIV-1Δ*vif*, while C2 and its point mutants strongly inhibited the virus (Fig. 3C). Immunoblot analysis of cell and viral lysates further confirmed that cellular expression and viral encapsidation of these variants were comparable (Fig. 3D). Finally, to test the *in vivo* DNA editing capacity, we did 3D-PCR analysis using C2, C2.DH-YG, and C2.RKYG-WEND variants. As presented in Fig. 3E, only HIV-1Δ*vif* particles produced in the presence of A3C chimera C2 and its mutant C2.DH-YG harbored viral DNA that was detected by PCR products at low-denaturation temperature and C2.RKYG-WEND behaved similarly to the vector control (Fig. 3E). Likewise, replacing RKYG with WEND in the smmA3C-like protein inhibited its antiviral activity (Figs. 4A and 4B) and deamination activity of HIV-1 genomes (Fig. 4C) as did the active site mutant E68A.

**Figure 3.**
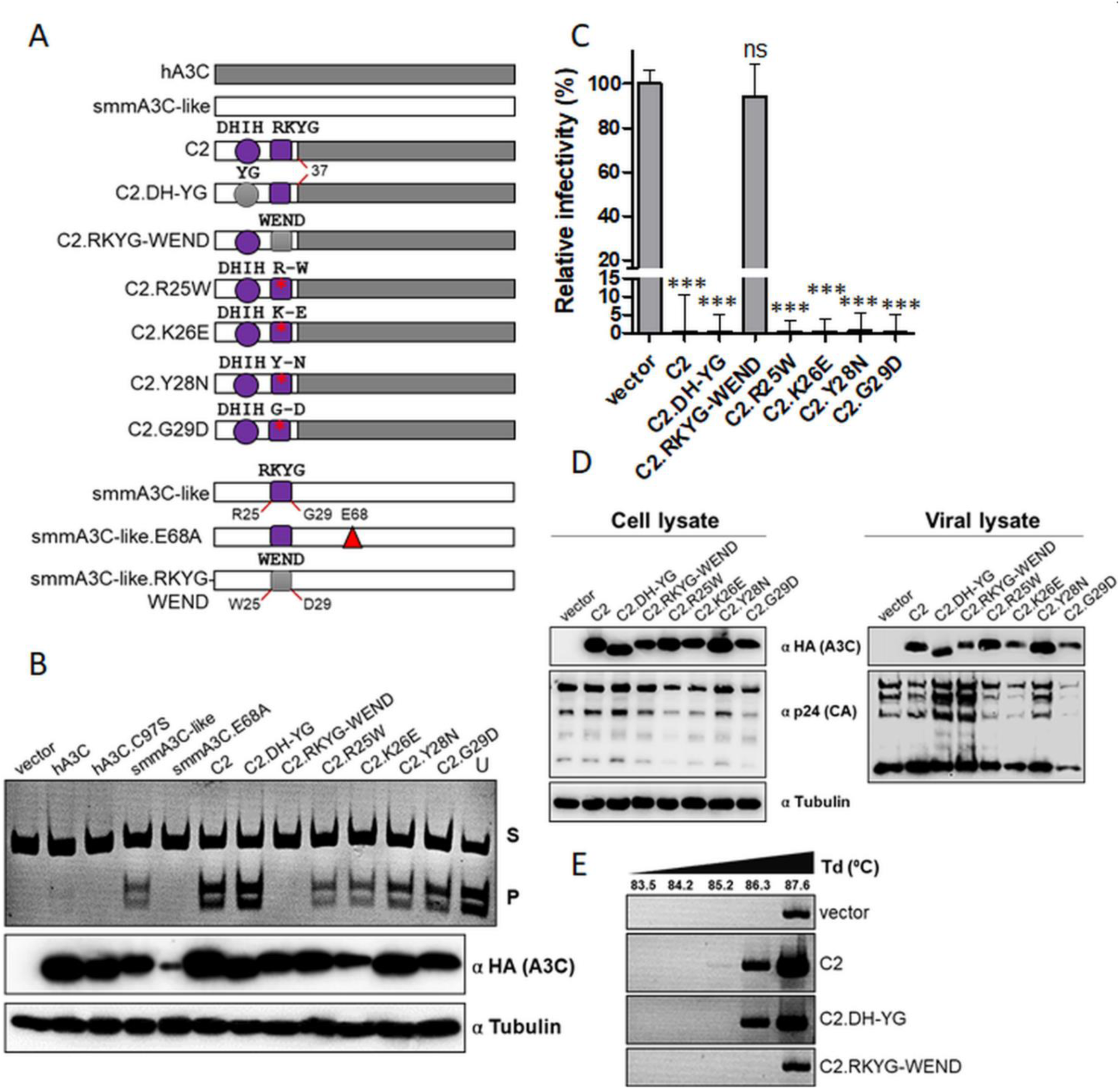
Identification of the A3Z2 domain mediating enhanced antiviral activity. (A) Illustration of chimera 2 (C2) and variants of C2 or smmA3C-like protein having amino acid exchanges in the DHIH (circle) or RKYG (square) motif. The red triangle denotes catalytic residue E68A mutation. Amino acid position (number) at the breakpoints of each chimera is indicated. Please see Suppl. Fig. S4 for more details about the sequence and structure of these motifs. (B) To examine the catalytic activity of chimeras C2 and its variants, *in vitro* deamination assays were performed using lysates of transfected cells. RNAse A-treatment was included; oligonucleotide containing uracil (U) instead of cytosine served as a marker to denote the migration of deaminated product after restriction enzyme cleavage. S-substrate, P-product. Immunoblot shows the amount of proteins produced in the transfected cells (lower panel). A3s were stained with anti-HA antibody and tubulin served as a loading control. “α” represents anti. (C) HIV-1Δ*vif* particles were produced with C2 and its variants or vector only. Infectivity of (RT-activity normalized) equal amounts of viruses, relative to the virus lacking any A3, was determined by quantification of luciferase activity in HEK293T cells. Values are means ± standard deviations (error bars) for three independent experiments. Unpaired t-tests were computed to determine whether differences between vector and each A3 protein reach the level of statistical significance. Asterisks represent statistically significant differences: ***, *p* < 0.0001; ns, not significant. (D) Amount of proteins in the cell lysate and viral encapsidation of C2 or its variants were determined by immunoblotting. C2/variants and HIV-1 capsids were stained with anti-HA and anti-p24 antibodies, respectively. Tubulin served as a loading control. (E) Quantification of hypermutations in viral DNA by 3D-PCR. HIV-1Δ*vif* particles produced in the presence of overexpressed C2, C2.DH-YG, C2.RKYG-WEND or vector controls were used to transduce HEK293T cells. Total DNA was extracted and a 714-bp fragment of reporter viral DNA was selectively amplified using 3D-PCR. T_d_ = denaturation temperature.

**Figure 4.**
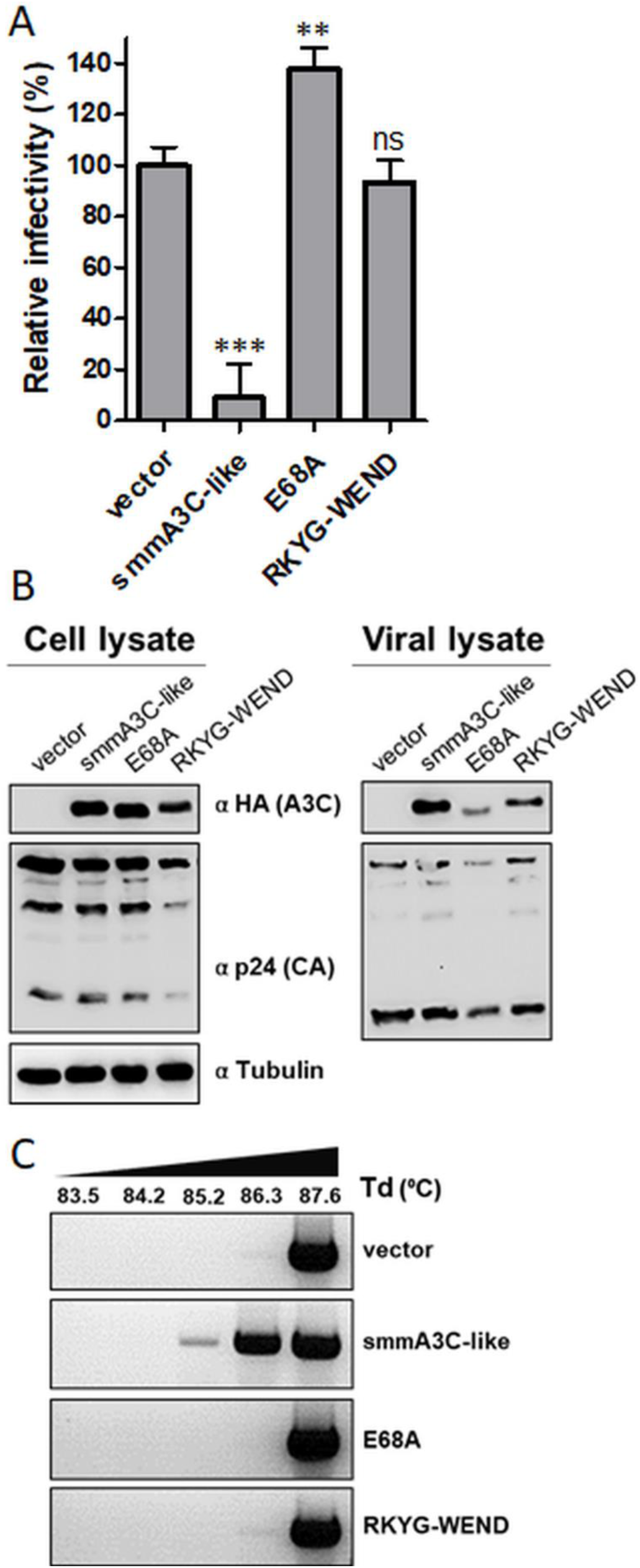
RKYG-WEND exchange in smmA3C-like protein abrogates its antiviral activity. (A) HIV-1Δ*vif* particles were produced with smmA3C-like protein, its mutants E68A (catalytically inactive), RKYG-WEND or vector only. Infectivity of (RT-activity normalized) equal amounts of viruses, relative to the virus lacking any A3, was determined by quantification of luciferase activity in HEK293T cells. (B) Immunoblot analyses were performed to quantify HA-tagged A3 proteins and viral p24 proteins in cellular and viral lysates using anti-HA and anti-p24 antibodies, respectively. Tubulin served as a loading control. “α” represents anti. (C) Quantification of hypermutation in viral DNA by 3D-PCR. HIV-1Δ*vif* particles produced in the presence of overexpressed smmA3C-like protein, its variants or vector control were used to transduce HEK293T cells. Total DNA was extracted and a 714-bp fragment of reporter viral DNA was selectively amplified using 3D-PCR. T_d_ = denaturation temperature.

### The WE-RK mutation in loop 1 of hA3C determines its strong deaminase-dependent antiviral function

Mutational changes of the RKYG motif to WEND residues in loop 1 of C2 and smmA3C-like protein resulted in complete loss of enzymatic functions and anti-HIV-1 activities (Figs. 3C and 4A). To identify the residues in hA3C that are critically required for the deaminase-dependent antiviral activity against HIV-1Δ*vif*, we modified the loop 1 of hA3C with ^25^WE^26^>>^25^RK^26^ and ^28^ND^29^>>^28^YG^29^ residues and compared their antiviral capacity (please see A3C alignment and ribbon diagram Suppl. Fig. S4). As controls, we included additional mutants such as a catalytically inactive Zn^2+^-coordinating C97 mutant, A3C.C97S [52], and the variants A3C.S61P [65] and A3C.S188I [69] (Suppl. Fig. S4A) exhibiting enhanced deaminase activity. Compared to wild-type hA3C, WE-RK greatly enhanced inhibition of HIV-1Δ*vif*, and the ND-YG variant behaved like wild-type A3C, while S61P and S188I have demonstrated only marginally increased HIV-1Δ*vif* restriction (Fig. 5A). Importantly, mutant A3C.C97S did not inhibit HIV-1Δ*vif* (Fig. 5A).

**Figure 5.**
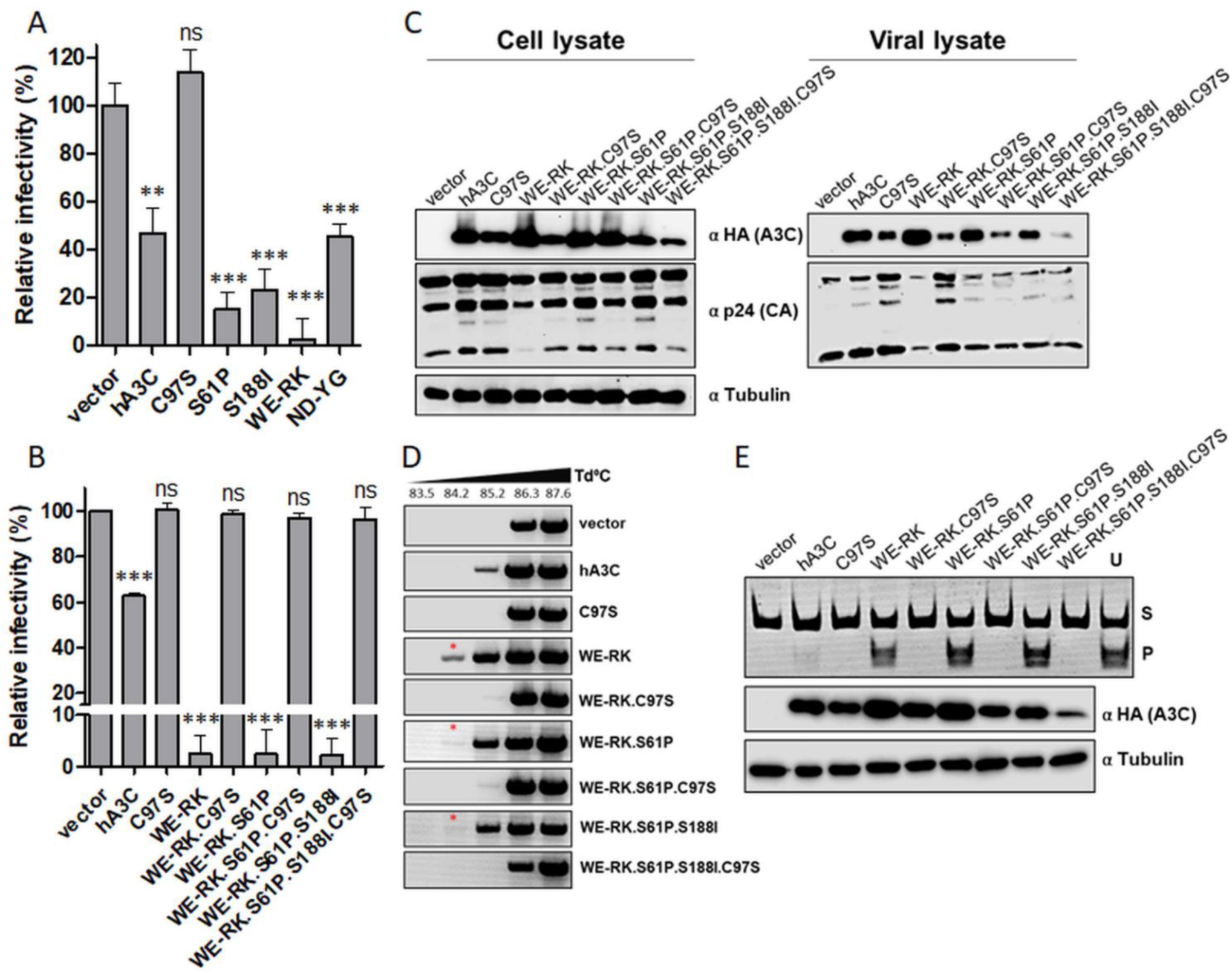
A3C gains deaminase-dependent anti-HIV-1 activity by a WE-RK change in loop 1. (A) HIV-1Δ*vif* particles were produced with hA3C, its mutants (C97S, S61P, S188I, WE-RK, ND-YG) or vector only. Infectivity of equal amounts of viruses (RT-activity normalized), relative to the virus lacking any A3C, was determined by quantification of luciferase activity in HEK293T cells. (B) HIV-1Δ*vif* particles were produced with hA3C, its variants such as, C97S, WE-RK, WE-RK.C97S, WE-RK.S61P, WE-RK.S61P.C97S, WE-RK.S61P.S188I, WE-RK.S61P.S188I.C97S or vector only. Infectivity of equal amounts of viruses (RT-activity normalized), relative to the virus lacking any A3C, was determined by quantification of luciferase activity in HEK293T cells. (C) Quantification of HA-tagged wild-type and mutant A3C proteins in both cellular and viral lysates by immunoblot analysis. A3s and HIV-1 capsids were stained with anti-HA and anti-p24 antibodies, respectively. Tubulin served as a loading control. “α” represents anti. (D) 3D-PCR: HIV-1Δ*vif* produced together with hA3C, its variants (as in Fig. 5B), or vector controls were used to transduce HEK293T cells. Total DNA was extracted and a 714-bp fragment of reporter viral DNA was selectively amplified using 3D-PCR. T_d_ = denaturation. Please see Suppl. Fig. S6 for the 3D-PCR data of mutants S61P and S188I. (E) *In vitro* deamination assays to examine the catalytic activity of A3C and its variants using lysates of cells that were previously transfected with respective expression plasmids (as in Fig. 5B). RNAse A-treatment was included; oligonucleotide containing uracil (U) instead of cytosine served as a marker to denote the migration of deaminated product after restriction enzyme cleavage. S-substrate, P-product. The two lower panels represent immunoblot analyses of expression levels of HA-tagged A3C and mutant proteins (α HA (A3C)) and tubulin (α tubulin) which was used as a loading control.

Next, we generated active site mutants to analyze if the antiviral activity of A3C.WE-RK is deamination-dependent. To achieve this, we introduced a C97S mutation in each of these constructs. Additionally, we compared the ancillary effect of mutants such as S61P [65] and S188I [69] by introducing these mutations in the WE-RK variant of A3C. As expected, the inhibitory activities of A3C.WE-RK, A3C.WE-RK.S61P, and A3C.WE-RK.S61P.S188I against HIV-1Δ*vif* were abolished by active site ablating mutation C97S, indicating the importance of the enzymatic activity of A3C (Fig. 5B). Introducing either the single mutation S61P or the double mutation S61P.S188I did not considerably change the action of A3C.WE-RK (Fig. 5B). Immunoblot analysis of cell and viral lysates demonstrated that hA3C and all mutants (except A3C.WE-RK.S61P.S188I.C97S mutant) expressed a comparable level of protein (Fig. 5C).

However, viral incorporation of A3C.C97S, A3C.WE-RK.C97S, A3C.WE-RK.S61P.C97S, and WE-RK.S61P.S188I.C97S was slightly decreased relative to that of wild-type and mutant proteins that do not contain the C97S mutation (Fig. 5C). Moreover, we confirmed the effects of these mutants on HIV-1Δ*vif* propagation by 3D-PCR (Fig. 5D) and deamination assay *in vitro* (Fig. 5E). In both assays, we found that the C97S mutation destroys the function of all A3C variants. Thus, we conclude that the loop 1-mediated enhanced activity of hA3C.WE-RK is dependent on catalytic deamination.

### The RK-WE mutation in loop 1 moderately reduces the antiviral activity of hA3F

The residues ^25^RK^26^ in loop 1 of smmA3C-like protein are derived from exon 5 of A3F gene in which exon 5 to 7 encoding A3F-CTD and conserved in primate A3F proteins (Suppl. Fig. S2). Various loops within A3F-CTD were recently investigated with respect to their role in substrate binding and enzyme function [70] but it was not possible to unravel the antiviral activity of A3F-CTD, mainly due to difficulties in expressing this domain in human cells as found earlier [65, 71]. hA3C and hA3F-CTD display 77% sequence similarity, reflecting a common evolutionary origin [6]. Importantly, the antiviral activity of hA3F is mediated by its CTD [72, 73]. To test the impact of RK residues in loop 1 of the hA3F-CTD, we compared the antiviral activity of hA3F with A3F.RK-WE against HIV-1Δ*vif*. hA3F and hA3F.RK-WE expressed similar amounts of protein and were equally encapsidated in HIV-1 particles (Fig. 6A). However, A3F.RK-WE exhibited an about two-fold decreased capacity to inhibit HIV-1Δ*vif* compared with wild-type A3F (Fig. 6B). Consequently, A3F.RK-WE showed decreased mutation efficiency compared with wild-type A3F (Figs. 6C and 6D), which was consistent with data presented in a recent report [70]. Thus, we conclude that loop 1 with its residues RK in CTD of A3F is important for hA3F’s enzymatic function.

**Figure 6.**
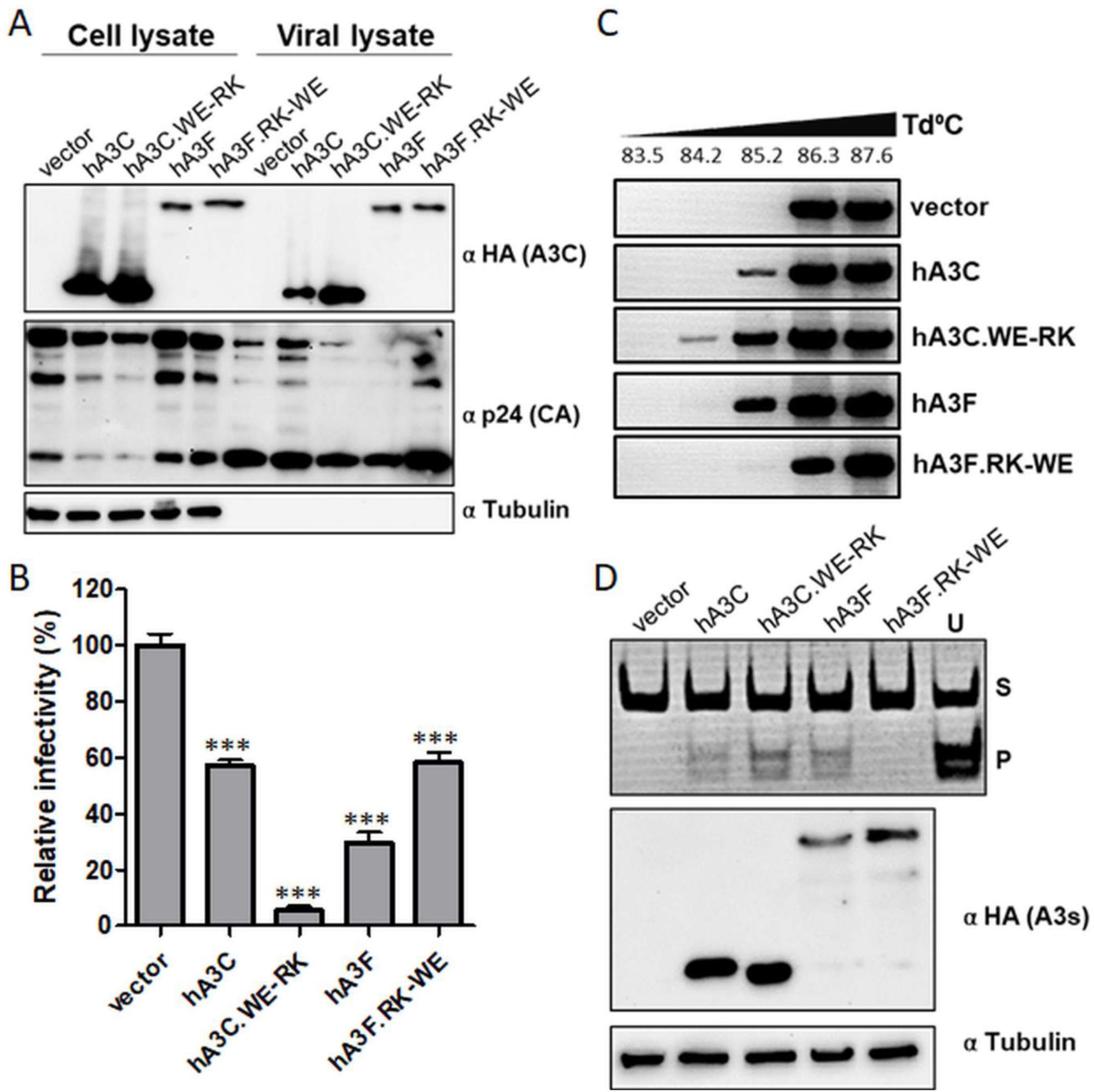
Mutations in loop 1 of A3F-CTD moderately affect the antiviral activity of A3F. (A) Immunoblot analyses were performed to quantify the amounts of HA-tagged wild-type hA3C and hA3F proteins and their loop 1 mutants in cell lysates and viral particles. HA-tagged A3s and HIV-1 capsid proteins were stained with anti-HA and anti-p24 antibodies, respectively. Tubulin served as a loading control. “α” represents anti. (B) Infectivity of equal amounts of HIV-1Δ*vif* viruses (RT-activity normalized) encapsidating hA3C, hA3F, or their loop 1 mutants relative to the virus lacking any A3 protein was determined by quantification of luciferase activity in transduced HEK293T cells. (C) 3D-PCR: HIV-1Δ*vif* produced together with hA3C, hA3F, and their loop 1 mutants or vector control were used to transduce HEK293T cells. Total DNA was extracted and a 714-bp fragment of reporter viral DNA was selectively amplified using 3D-PCR. T_d_ = denaturation temperature. (D) *In vitro* deamination assay to examine the catalytic activity of hA3C, hA3F, and their loop variants was performed using lysates of cells that were transfected with the respective A3 expression plasmids. RNAse A-treatment was included; oligonucleotide containing uracil (U) instead of cytosine served as a marker to denote the migration of the deaminated products after restriction enzyme cleavage. S-substrate, P-product. The two lower panels represent immunoblot analyses of expression levels of HA-tagged A3C, A3F and mutant proteins (α HA (A3s)) and tubulin (α tubulin) which was used as a loading control.

### Inhibition of LINE-1 retrotransposition by A3C variants

Since A3C and A3F restrict endogenous LINE-1 (L1) retrotransposition activity by 40-75% and 66-85% [46,56,74,75], respectively, we set out to elucidate how the WE and the RK residues in loop 1 of both hA3C and hA3F, respectively, affect the L1 inhibiting activity. To this end, we quantified the L1-inhibiting effect of human wild-type A3A, A3C, and A3F proteins and their mutants hA3C.WE-RK, hA3C.WE-RK.S61P, and hA3F.RK-WE by applying a dual-luciferase retrotransposition reporter assay [76]. In this cell culture-based assay, the firefly luciferase gene is used as the reporter for L1 retrotransposition and the Renilla luciferase gene is encoded on the same plasmid for transfection normalization (Fig. 7A). Consistent with previous reports, overexpression of hA3A, hA3C, and hA3F resulted in inhibition of L1 reporter retrotransposition by approximately 94%, 68%, and 56%, respectively (Fig. 7B). The mutant hA3C.WE-RK displayed an increased L1-restricting effect (from 56% to ∼96%), and the introduction of the additional mutation hA3C.WE-RK.S61P did not further increase the ability of the enzyme to restrict L1 mobilization (Fig. 7B). Notably, hA3F and the mutant hA3F.RK-WE exhibited a comparable level of L1 restriction, indicating that regions other than loop 1 of A3F-CTD and, probably, the NTD (N-terminal domain) of hA3F are involved in L1 restriction (Fig. 7B). Immunoblot analysis of cell lysates of co-transfected HeLa-HA cells demonstrated comparable expression of the L1 reporter and HA-tagged A3- and A3 mutant proteins (Suppl. Fig. S7). These findings indicate that the WE-RK mutation in hA3C enhances its L1 inhibiting activity. Based on the observed antiviral activity and the L1 restricting effect of hA3C.WE-RK on L1, we hypothesize that the introduction of these positively charged residues in hA3C significantly fosters its interaction with nucleic acids, which was recently reported to mediate its L1 inhibiting activity [56].

**Figure 7.**
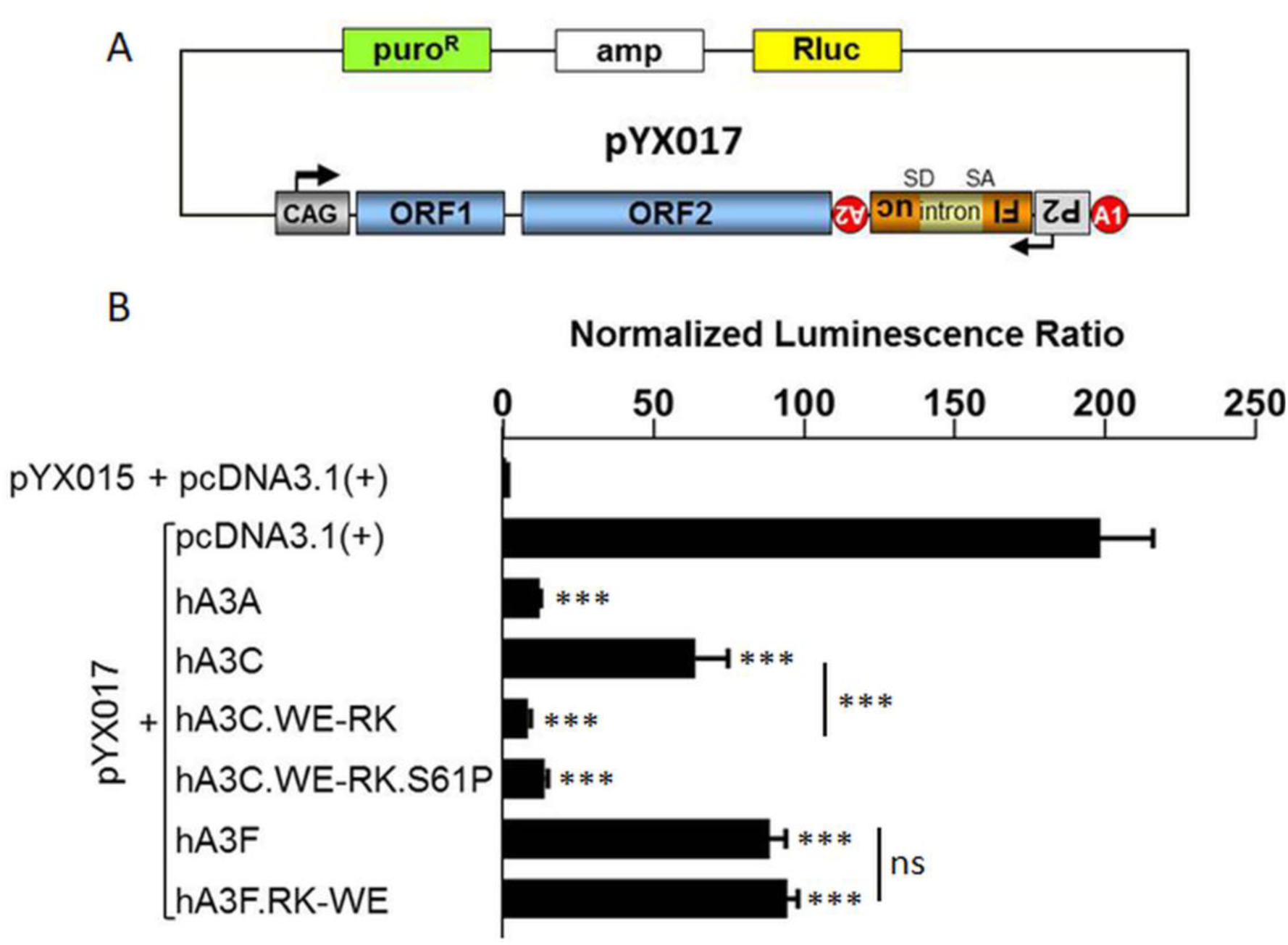
Expression of the hA3C.WE-RK variant enhances A3C-mediated L1 restriction significantly. Dual-luciferase reporter assay to evaluate the effect of wild-type and mutant A3 proteins on L1 retrotransposition activity. (A) Schematic of the L1 retrotransposition reporter construct pYX017 [76]. The L1_Rp_ reporter element is under transcriptional control of the CAG promoter and a polyadenylation signal (A1) at its 3‘end. The firefly luciferase (Fluc) cassette has its own promoter (P2) and polyadenylation signal (A2), is expressed from the antisense strand relative to the CAG promoter, and interrupted by an intron (with splice donor [SD] and splice acceptor [SA]) in the transcriptional orientation of the L1 reporter element. (B) Effect of wild-type and mutant A3 proteins on L1 retrotransposition activity indicated by normalized luminescence ratio (NLR). NLR indicating retrotransposition activity observed after cotransfection of pYX015 and empty pcDNA3.1 (+) expression plasmid was set as 1. Error bars indicate standard deviation (N=4). The protein expression level of A3s is presented in Suppl. Fig. S7.

### The positively charged residues R25 and K26 in A3C-C2 form salt-bridges with the backbone of the ssDNA

The structural model of hA3C variant C2 binding to ssDNA, which is based on the ssDNA-bound crystal structure of A3A, shows a cytidine residue in the active center of hA3C-C2 (Fig. 8A). However, the ssDNA fragment, which was co-crystallized with hA3A, is too short to interact with residues 25, 26, 28, and 29, which differ between hA3C WT and the C2 variant. Hence, this binding mode model cannot explain why C2 has a higher cytidine deaminase activity than hA3C WT. To assess the binding to a longer ssDNA fragment, we generated a complex model of ssDNA bound to the NTD of rhesus macaque A3G (rhA3G) [77], similar to the ssDNA-bound A3F-CTD model built previously [78], and aligned the crystal structure of hA3C WT and the model of C2 to this complex (Figs. 8B, 8C, and 8D). The positively charged residues R25 and K26 in C2 form salt-bridges with the backbone of the ssDNA (Fig. 8D) in contrast to hA3C WT (Fig. 8C). Additionally, Y28 of C2 can form π-π-stacking interactions with the aromatic DNA bases (Fig. 8D). Thus, these three residues can form stronger interactions with ssDNA in C2 than their counterparts in hA3C. This finding may explain the enhanced cytidine deaminase activity of C2 compared to hA3C.

**Figure 8.**
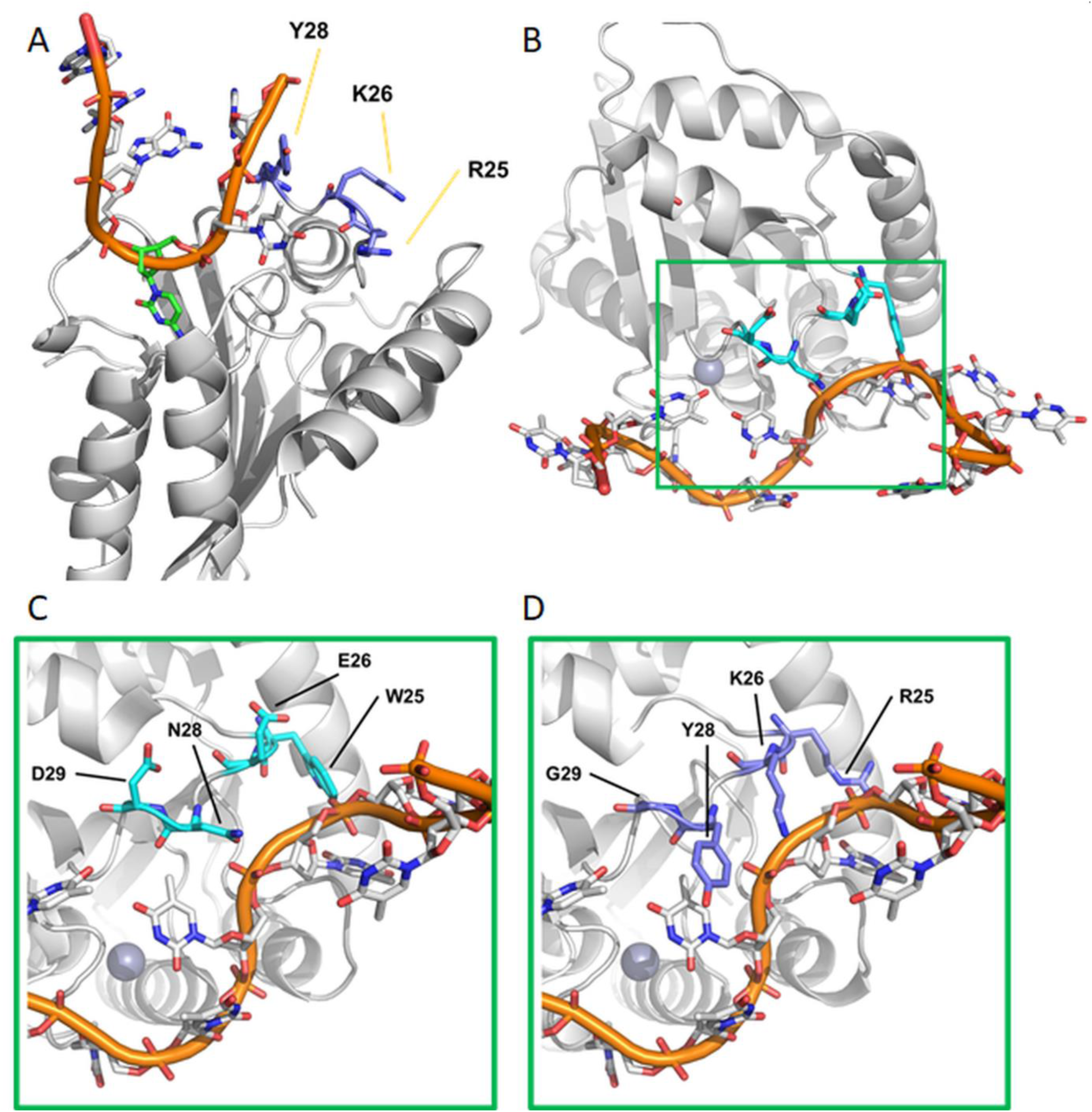
Model of A3C.WE-RK and ssDNA interaction. (A) Binding mode model of ssDNA (orange) to hA3C variant C2 based on hA3A. The side chains of the residues that are different between the hA3C WT and the C2 variant are shown in dark blue and labelled. The cytidine in the active center of C2 is shown in green. (B) Binding mode model of ssDNA (orange) to hA3C WT based on hA3F-CTD and rhA3G-NTD. Magnifications of the active center (green box) are shown at the bottom for hA3C WT (C) and C2 (D). The side chains of residues in the active center that differ between hA3C WT and the C2 variant are shown in cyan and dark blue, respectively. The Zn^2+^ ion in the active center is shown as a sphere. Ongoing from hA3C WT to the C2 variant, the interface changes from being negatively to being positively charged. The flexible arginine and lysine side chains in the C2 variant can interact with the negatively charged backbone of ssDNA (panel D), stabilizing this interaction.

Furthermore, we performed five replicas of molecular dynamics (MD) simulations of 2 µs length each for hA3C, C2, and hA3C.S61P.S188I to assess the structural impact of the substitutions. The root mean square fluctuations (RMSF), which describe atomic mobilities during the MD simulations, show distinct differences between the variants in the putative DNA-binding regions of the proteins: the RMSF of C2 and hA3C.S61P.S188I are up to 2 Å larger compared to hA3C WT in the regions carrying the substitutions (residues 21-32 for C2 and residues 55-67 for hA3C.S61P.S188I) (Suppl. Fig. S8). This effect is specifically related to the respective substitutions, as no change in RMSF occurs for a variant in a region where it is identical to A3C WT. The increased movement of ssDNA-binding residues might improve the sliding of C2 and hA3C.S61P.S188I along the ssDNA, owing to more transient interactions with the ssDNA backbone. Conversely, the RMSF of loop 7 is up to 1 Å lower in both the C2 and hA3C.S61P.S188I variants compared to the hA3C WT (Suppl. Fig. S8).

### WE-RK mutation in the loop 1 of hA3C enhances the interaction with ssDNA

To validate our structural modeling analysis (Fig. 8), and to address if the interaction of hA3C and hA3C.WE-RK with the substrate ssDNA was differentially affected, we performed electrophoretic mobility shift assays (EMSA) using hA3C-GST (A3C fused to glutathione S-transferase, GST) and hA3C.WE-RK-GST purified from HEK293T cells (Fig. 9A). As a probe, we used a biotin-labeled ssDNA oligonucleotide that harbors a TTCA motif in its central region [65, 79]. Because hA3C-GST is known to form a stable DNA-protein complex when the protein concentration reaches ≥ 20 nM ([65] and data not shown), we decreased the amount of A3C and its mutant protein to specifically test their inherent DNA binding capacity. In a titration experiment with concentrations ranging from 2 to 8 nM in steps of 2 nM of hA3C-GST and hA3C.WE-RK-GST purified protein, we detected a clear trend in the formation of DNA–protein complexes for hA3C-GST and hA3C.WE-RK-GST (Fig. 9B). Intriguingly, DNA-protein complexes of hA3C.WE-RK-GST started appearing at the lowest protein concentration used (2 nM), while hA3C-GST-DNA complexes were detected at protein concentration ≥ 6 nM. The top-shifted complexes were formed only with hA3C.WE-RK-GST and not with hA3C-GST. To confirm the specificity of the DNA–protein complexes, we competed for the reaction with unlabeled DNA carrying the same nucleotide sequence as the used probe in 500-fold excess relative to that probe. The addition of the competitor DNA to the sample containing the maximum (8 nM) amount of A3C protein, efficiently disrupted the protein-DNA complex formation. Together, data from structural modeling and EMSA experiments allowed us to conclude that the two amino acid-change in loop 1 of A3C boosts the ssDNA binding capacity of A3C. Importantly, the GST moiety did not affect the binding ([65] and data not shown).

**Figure 9.**
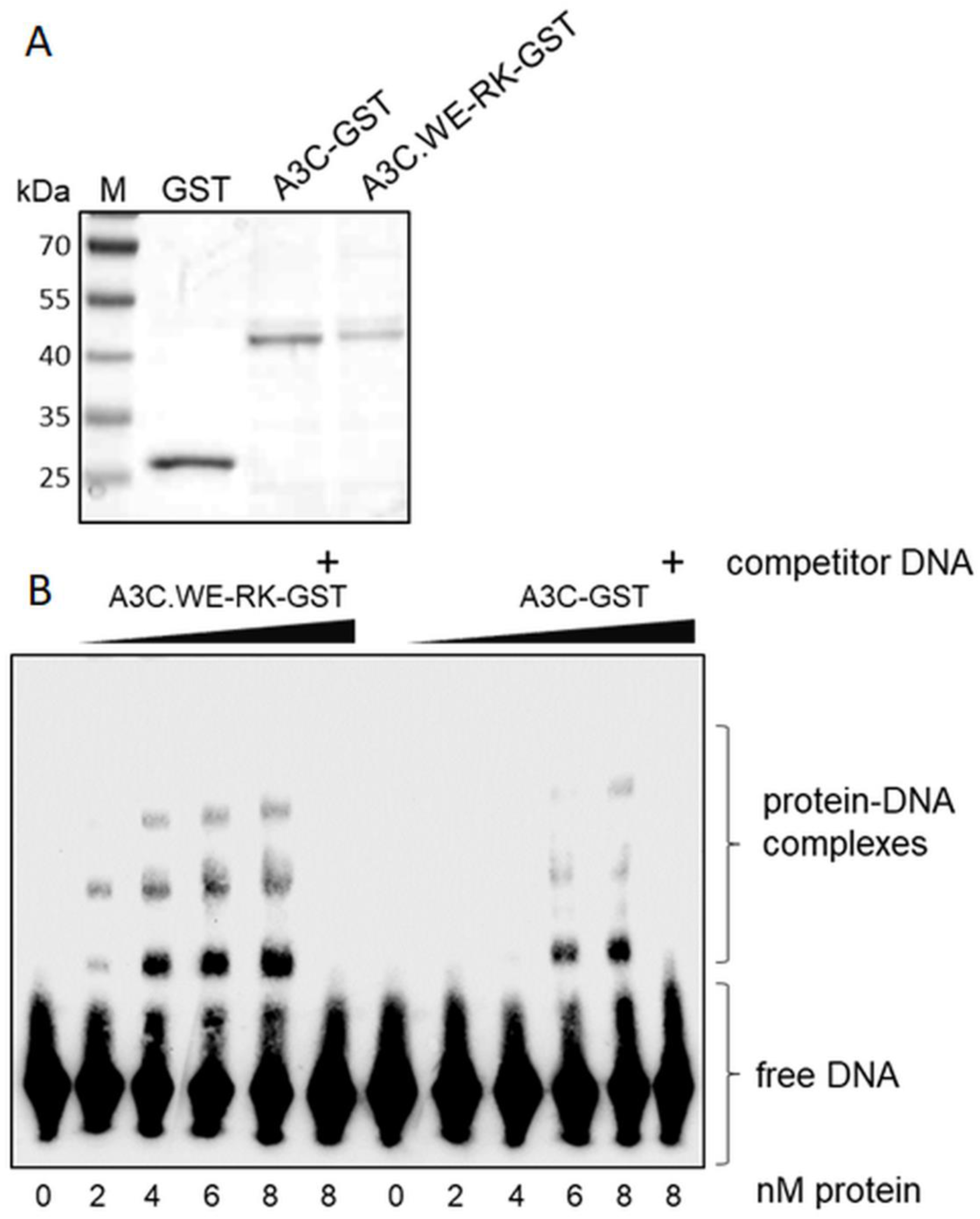
hA3C.WE-RK strongly interacts with ssDNA. (A) The purity of the recombinantly produced and affinity-purified proteins GST, A3C-GST, and A3C.WE-RK-GST was demonstrated by SDS-PAGE and subsequent Coomassie blue staining of the gel. The prestained protein ladder (M) indicates molecular mass. (B) EMSA with GST-tagged hA3C.WE-RK-GST and A3C-GST produced in HEK293T cells was performed with 30-nt ssDNA target DNA labelled with 3′-labelled biotin. Indicated amounts of protein (at the bottom of the blot, in nm) were titrated with 20 fmol of DNA. Presence of competitor DNA (unlabeled 80-nt DNA used in deamination assay, 500-fold molar excess added) used to demonstrate the specific binding of the protein to DNA being causative for the shift.

### Evolution of A3Z2 loop 1 regions in primates

We performed a phylogenetic reconstruction for the A3Z2 domains in primates, using the A3Z2 sequences in the northern tree shrew as outgroup. Because in primates A3D and the A3F contain two Z2 domains, we analyzed at the A3Z2 domain level (N- and C-terminal Z2) (Fig. 10A). Our results show that the A3Z2 domains underwent independent duplication in the two sister taxa, tree shrews and primates: the three A3Z2 tree shrew sequences constitute a clear outgroup to all primate A3Z2 sequences. We identified a sharp clustering of the A3D-NTD and A3F-NTD on the one hand and of A3C, A3D-CTD, and A3F-CTD on the other hand. As to New World monkeys (Platyrrhini), we could only confidently retrieve A3C sequences from the white-faced sapajou *Cebus capucinus* and from the Ma’s night monkey *Aotus nancymaae.* These sequences from A3C New World monkeys were basal to all Catarrhini (Old World monkeys and apes) A3C, A3D-CTD and A3F-CTD sequences, suggesting that the two gene duplications leading to the extant organization of A3C, A3D, and A3F occurred after the Platyrrhini/Catarrhini split 43.2 Mya (41.0 - 45.7 Mya) and before the Cercopithecoidea/Hominoidea split 29.44 Mya (27.95 - 31.35 Mya). The results show a tangled distribution within the A3D-NTD and A3F-NTD clade, and within the A3D-CTD and A3F-CTD clade. These confusing relationships are more obvious when comparing an unconstrained Z2 tree with a tree in which monophyly of the large six clades identified was enforced. Conversely, Catarrhini A3C sequences form a monophyletic taxon, and this A3C gene tree essentially adheres to the corresponding species tree (Fig. 10B). Focusing on the nodes that we could identify with confidence, we performed ancestral phylogenetic inference of the most likely amino acid sequence for the A3 loop 1 as well as consensus analysis of the extant sequences. Our results recover the well-conserved aromatic stacking stretch F[FY]FXF characteristic of all A3s. In the A3C, A3D-CTD, and A3F-CTD clade, we identified a motif with divergent evolution flanked by conserved small hydrophobic amino acids. The most likely ancestral form is the amino acid motif LRKA, which is also the form present in extant New World monkeys A3C and the most common in extant A3F-CTD; in the extant A3D-CTD the Arg residue is less conserved in L[RLQ][KT]A; and strikingly, in the ancestor of Catarrhini A3C, this motif changed to LWEA. Only subsequently, and exclusively in the Chlorocebus lineage, this change was partly reverted to LREA by a transition TGG>CGG. This reversion should have occurred after the divergence within Cercopithecinae, around 13.7 Mya (10.7 - 16.6 Mya).

**Figure 10A.**
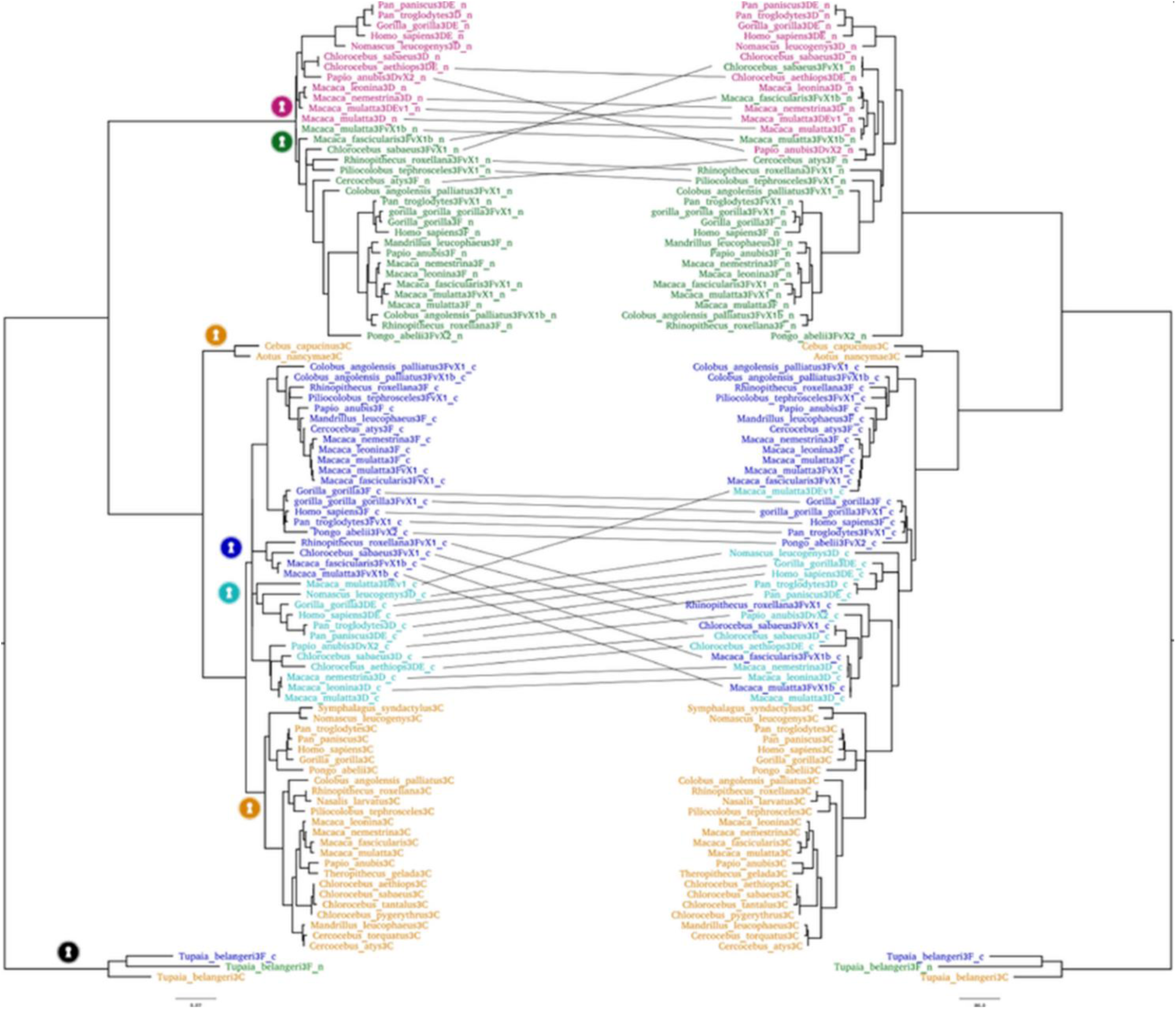
Evolution of A3C/Z2. Tanglegram showing the maximum-likelihood amino acid-based reconstructions of the A3 Z2 sequences in primates. The A3D and A3F proteins, containing each two Z2 domains, were split into N- and C-termini. The analyses were performed at the Z2-domain level, labelled as follows: A3D-NTD, red; A3F-NTD, green; A3D-CTD, light blue; A3F-CTD, dark blue; A3C, orange. The A3Z2 sequences of the northern tree shrew *Tupaia belangeri* were used as an outgroup. On the right, the phylogenetic reconstruction was built without any topological restriction. On the left, monophyly of the different Z2 domains was enforced, as indicated with the keyhole symbols of the corresponding color. Sequences with a significantly different position in the two trees are connected by a line. All A3C sequences are consistently monophyletic, and their phylogenetic relationships in the gene tree adhere to those of the species tree. Conversely, for Old World monkeys, sequences identified as A3D-CTD are in certain cases closest relatives of A3F-CTD, and *vice versa*. Such anomalies could have a methodological basis, such as poor annotation or overlooked chimeric amplification of closely related sequences into a single amplicon. Alternatively, they could be related to a genuine biological phenomenon of interlocus gene conversion, in which one allele (or one genomic stretch) is replaced by a paralogous allele with which it shares a high sequence identity. In this case, the NTD and the CTD of A3D and A3F share respectively very recent common ancestors.

**Figure 10B.**
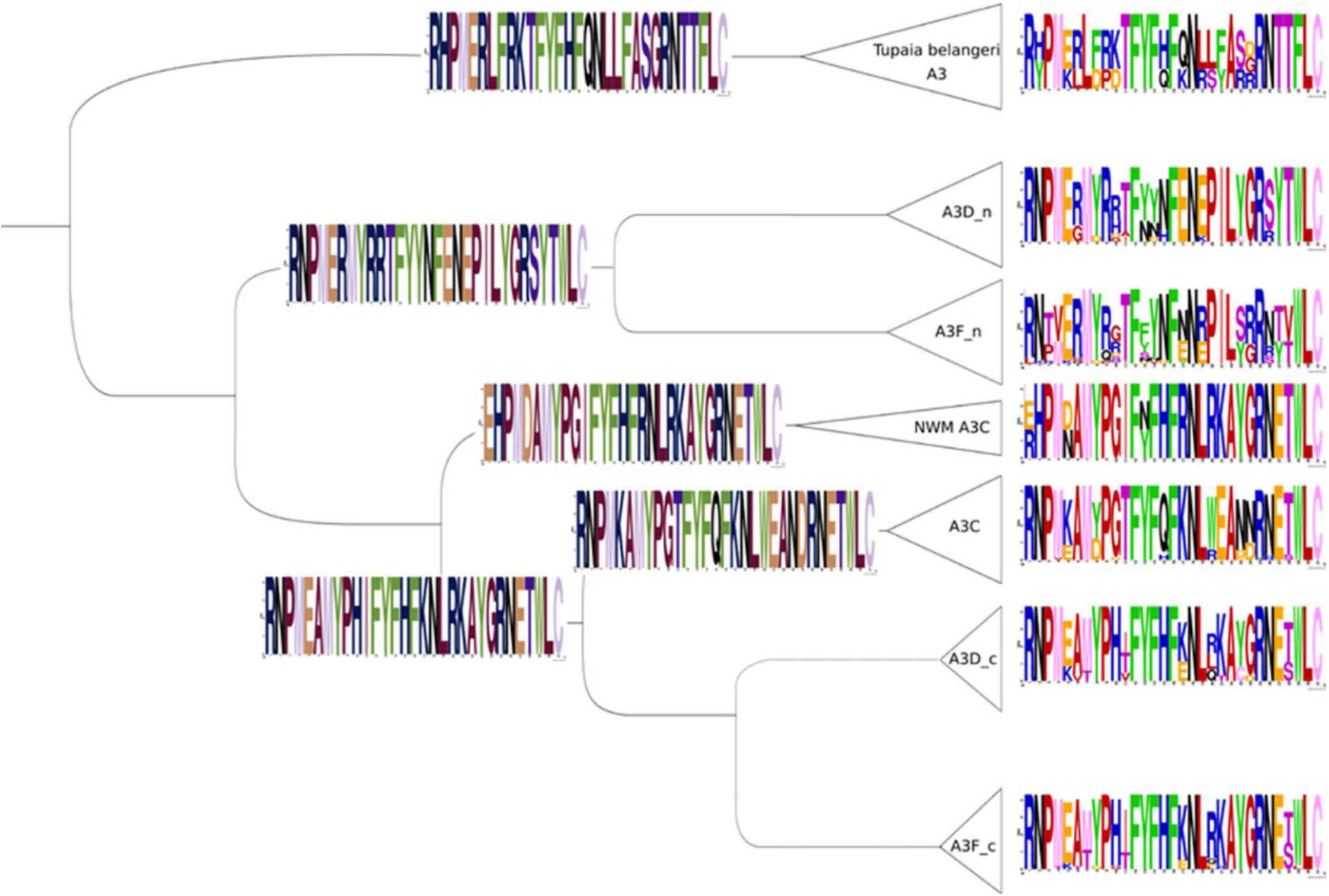
Evolution of A3C/Z2. Phylogenetic relationships between A3 Z2 sequences in primates, using A3 Z2 sequences in the northern tree shrew *Tupaia belangeri* genome as an outgroup (see Suppl. Fig. S10 for evolutionary relationships of analyzed species). The A3D and A3F proteins, containing each two Z2 domains, were split into NTD and CTD. The A3C sequences in the New World Monkeys (A3C_NWM) are monophyletic and constitute the sister taxa for A3C, A3D-CTD, and A3F-CTD. For each A3 Z2 sequence cluster, a sequence logo focused on loop 1 of the extant sequences is shown on the right. For the ancestor nodes that can be inferred with confidence, the most likely amino acid sequence of the ancestral state is provided. The last common ancestor of A3C, A3D-CTD and A3F-CTD likely presented a RKAYG sequence in the protein loop 1, and this sequence has been maintained in New World Monkeys’ A3Cs. However, this sequence stretch underwent a strong selective pressure and became the strongly divergent WEA[ND] [ND] in present-day A3Cs. Subsequently, and exclusively in A3C in the Chlorocebus genus, a secondary W>R reversion occurred. The ancestral sequence was essentially maintained [RQ]KAYG in A3F-CTD, while it was largely eroded [RLQ]KA [YG] [CD] in A3C.

## DISCUSSION

Compared to the studies conducted over the past decade on the potent HIV-1 restriction factors A3G and A3F, investigations on A3C are very limited. Only a few recent studies have addressed the catalytic activity and substrate binding capacity of A3C [65,69,80]. While the previously characterized hA3C mutants S61P and S188I boost the catalytic activity of the enzyme to a certain level, none of these mutations are decisive because they do not reduce the HIV-1Δ*vif* infectivity to any level accomplished by A3G nor do they directly partake in catalytic activity [65,69,80]. Because our repeated attempts to express A3F-CTD in human cells were not successful ([65] and Suppl. Fig. S3), we assayed A3C proteins from different Old World monkey species. Due to the high level of nucleotide sequence identity between the A3 paralogs in the sooty mangabey monkey genome, we unintentionally generated the smmA3C-like protein with superior anti-HIV-1 and enzymatic activity. We have identified the key role of two positively-charged residues in loop 1 of the smmA3C-like protein (and of the hA3F-CTD), namely R25 and K26 in the RKYG motif. Replacing RKYG of A3C chimera C2 or smmA3C-like protein by the WEND (form of this motif in hA3C) abolished both their anti-HIV-1 and catalytic activity. Notably, the converse strategy of introducing the substitution WE-RK in the loop 1 of hA3C rendered hA3C.WE-RK a potent, deaminase-dependent, anti-HIV-1 enzyme. Consistent with these observations, our EMSA data clearly demonstrate that residues in the loop 1 of A3C regulate protein-DNA interaction and we postulate that this interaction is causative for the enhanced deamination activity and enhanced anti-HIV and L1 activity. A similar model was discussed by Solomon and coworkers, which demonstrated that loop 1 residues of hA3G-CTD-2K3A-E259A (a catalytically inactive form of A3G-CTD) strongly interact with substrate ssDNA and that this distinguishes catalytic binding from non-catalytic binding [81]. Interestingly, loop 1 of A3A was found to be important for substrate specificity but not for substrate binding affinity [82], while loop 1 of A3H especially residue R26, plays a triple role for RNA binding, DNA substrate recognition, and catalytic activity likely by positioning the DNA substrate in the active site for effective catalysis [83]. In accordance with this, our study claims that ^25^RK^26^ substitution in loop 1 of A3C provides the microenvironment that drives the flexibility in substrate binding and enzymatic activity.

The binding model developed here rationalizes how A3C variant C2 can interact with the negatively charged backbone of ssDNA via the positively charged loop 1 side chains of R25 and K26 (Fig. 8D). Like our modeling strategy, Fang *et al*. [78] used their binding mode model of A3F-CD2 with ssDNA to identify residues in the A3G-CTD important for ssDNA binding. Furthermore, the increased mobility of DNA binding regions carrying the substitutions in C2 and hA3C-S61P.S188I, respectively, compared to hA3C (Suppl. Fig. S8) suggests that C2 and hA3C-S61P.S188I can better slide along the ssDNA than hA3C: The higher mobility of the residues may allow them to adapt more quickly to the passing ssDNA, which, paired with likely stronger interactions with the backbone of the ssDNA, may explain the increased deaminase activity. In addition, loop 7 exhibits a decreased mobility in both C2 and hA3C-S61P.S188I compared to hA3C, which was shown to be a predictor for higher deaminase activity, DNA binding, and substrate specificity of A3G and A3F, and reported to be also relevant for antiviral activity of A3B and A3D [73,84–86].

Unexpectedly, our experiments also demonstrated that LINE-1 restriction by A3C which was reported earlier to be deaminase-independent [56], is enhanced after expression of the A3C.WE-RK variant. These data suggest that the reported RNA-dependent physical interaction between L1 ORF1p and A3C dimers might be mediated by A3C loop 1, is partly dependent on the two amino acids W25 and E26 and enhanced by the R25 and K26 substitutions. However, L1 inhibition by A3F was not significantly altered by the A3F.RK-WE mutations, clearly indicating that in A3F other regions (and NTD) are likely to be relevant for L1 restriction.

Because selection likely had to balance between anti-viral/anti-L1 activity and genotoxicity of A3 proteins, we wanted to characterize loop 1 residues during the evolution of closely related A3Z2 proteins such as A3C, A3D CTD and A3F CTD in primates. In the most recent common ancestor of these enzymes, before the split Catarrhini-Platyrrhini some 43 Mya, the sequence of this motif in loop 1 is LRKAYG. In New World Monkeys, the A3C genes were not duplicated and are basal to the three sister clades of Catarrhini A3C, A3D-CTD, and A3F-CTD. In extant A3C sequences in New World monkeys, the loop 1 motif has notably remained unchanged and reads LRKAYG. In Catarrhini, on the contrary, the ancestral A3C sequence underwent two rapid rounds of duplication that occurred after the split with the ancestor of Platyrrhini, and before the split between the ancestors of Cercopithecoidea and Hominoidea, some 29 Mya. In extant A3F-CTD sequences, the consensus form of the loop 1 remains LRKAYG, albeit with certain variability of the Arg residue to be exchanged by other positively charged amino acids. In extant A3D-CTD enzymes, this motif has undergone erosion, is more variable and reads L[RLQ][KT]A[YC]G. Interestingly, loop 1 in A3C has experienced rapid and swift selective pressure to exchange the positively charged RK amino acids by the largely divergent chemistry of WE, yielding LWEAYG. This selective sweep occurred very rapidly, as this is the fixed form in all Catarrhini. Notoriously, and exclusively in the Chlorocebus lineage, this amino acid substitution was partly reverted to LREAYG, which is the conserved sequence in the four Chlorocebus A3C entries available.

Overall, our results suggest that the two duplication events that generated the extant A3C, A3D-CTD, and A3F-CTD sequences in Catarrhines, released the selective pressure on two of the daughter enzymes allowing them to explore the sequence space and to evolve via sub/neofunctionalisation, as proposed for Ohno’s in-paralogs [87]. Thus, the A3F-CTD form of the loop 1 diverged little from the ancestral chemistry and possibly maintained the ancestral function, while the release in conservation pressure on A3D-CTD allowed the enzyme loop 1 to accumulate mutations and diverge from the ancestral state. In turn, A3C was rapidly engaged into a distinct evolutionary pathway, which is unique due to the highly divergent chemistry of loop 1 but also because A3C is the only A3Z2 monodomain enzyme of the A3 family.

In conclusion, we postulate that the loop 1 region of A3s might have a conserved role in anchoring ssDNA substrate for efficient catalysis and that hA3C’s weak deamination and anti-HIV-1 activity might have been the result of losing DNA interactions in loop 1 during its evolution. It is thus possible that genes encoding A3C proteins with loop 1 residues with a higher ssDNA affinity were too genotoxic to benefit its host by superior anti-viral and anti-L1 activity.

## MATERIALS AND METHODS

### Cell culture

HEK293T cells were maintained in Dulbecco’s high-glucose modified Eagle’s medium (DMEM) (Biochrom, Berlin, Germany), supplemented with 10% fetal bovine serum (FBS), 2 mM L-glutamine, 50 units/ml penicillin, and 50 µg/ml streptomycin at 37^°^C in a humidified atmosphere of 5% CO_2_. Similarly, HeLa-HA cells [88] were cultured in DMEM with 10% FCS (Biowest, Nuaillé, France), 2mM L-glutamine and 20 U/ml penicillin/streptomycin (Gibco, Schwerte, Germany).

### Plasmids

The HIV-1 packaging plasmid pMDLg/pRRE encodes *gag-pol*, and the pRSV-Rev for the HIV-1 *rev* [89]. The HIV-1 vector pSIN.PPT.CMV.Luc.IRES.GFP expresses the firefly luciferase and GFP reported previously [90]. HIV-1 based viral vectors were pseudotyped using the pMD.G plasmid that encodes the glycoprotein of VSV (VSV-G). SIVagm luciferase vector system was described before [31]. All APOBEC3 constructs described here were cloned in pcDNA3.1 (+) with a C-terminal hemagglutinin (HA) tag. The smmA3C-like expression plasmid was generated by exon assembly from the genomic DNA of white-crowned mangabey (*Cercocebus torquatus lunulatus*), and the cloning strategy for smmA3C-like and the chimeras of hA3C/smmA3C-like plasmid construction was recently described [64]. The expression vector for A3G-HA was generously provided by Nathaniel R. Landau. Expression constructs hA3C, rhA3C, cpzA3C, agmA3C and A3C point mutant A3C.C97S were described before [52,55,65]. smmA3C-like with C-terminal V5 tag was cloned using following primers forward 5’-EcoRI-ATGAATTCGCCACCATGAATCCACAGATCAGAAAC and reverse 5’-NotI-ATGCGGCCGCCACTCGAGAATCTCCTGTAGGCGTC.

Various point mutants hA3C.WE-RK, hA3C.ND-YG, hA3C.WE-RK.C97S, hA3C.WE-RK.S61P, hA3C.WE-RK.S61P.C97S, hA3C.WE-RK.S61P.S188I, hA3C.WE-RK.S61P.S188I.C97S, hA3F.RK-WE, smmA3C-like.E68A were generated by using site-directed mutagenesis. Similarly, single or multiple amino acid changes were made in expression vectors to produce chimera 2 mutants (C2.DH-YG, C2.RKYG-WEND, C2.R25W, C2.K26E, C2.Y28N, and C2.G29D) and smmA3C-like.RKYG-WEND. To clone C-terminal GST-tagged hA3C, hA3C.WE-RK, the ORFs were inserted between the restriction sites HindIII and XbaI in the mammalian expression construct pK-GST mammalian expression vector [91]. Individual exons of authentic smmA3C and smmA3F and smmA3F-like genes exons were amplified and cloned in pcDNA3.1. All the primer sequences are listed in Suppl. table 1.

### Virus production and isolation

HEK293T cells were transiently transfected using Lipofectamine LTX and Plus reagent (Invitrogen, Karlsruhe, Germany) with an appropriate combination of HIV-1 viral vectors (600 ng pMDLg/pRRE, 600 ng pSIN.PPT.CMV.Luc.IRES.GFP, 250 ng pRSV-Rev, 150 ng pMD.G with 600 ng A3 plasmid or replaced by pcDNA3.1, unless otherwise mentioned) or SIVagm vectors (1400 ng pSIVTan-LucΔ*vif*, 150 ng pMD.G with 600 ng A3 plasmid) in 6 well plate. 48 h post-transfection, virion containing supernatants were collected and for isolation of virions, concentrated by layering on 20% sucrose cushion and centrifuged for 4 h at 14,800 rpm. Viral particles were re-suspended in mild lysis buffer (50 mM Tris (pH 8), 1 mM PMSF, 10% glycerol, 0.8% NP-40, 150 mM NaCl and 1X complete protease inhibitor).

### Luciferase-based infectivity assay

HIV-1 luciferase reporter viruses were used to transduce HEK293T cells. Prior infection, the amount of reverse transcriptase (RT) in the viral particles was determined by RT assay using Cavidi HS kit Lenti RT (Cavidi Tech, Uppsala, Sweden). Normalized RT amount equivalent viral supernatants were transduced. 48 h later, luciferase activity was measured using SteadyliteHTS luciferase reagent substrate (Perkin Elmer, Rodgau, Germany) in black 96-well plates on a Berthold MicroLumat Plus luminometer (Berthold Detection Systems, Pforzheim, Germany). Transductions were done in triplicate and at least three independent experiments were performed.

### Immunoblot analyses

Transfected HEK293T cells were washed with phosphate-buffered saline (PBS) and lysed in radioimmunoprecipitation assay buffer (RIPA, 25 mM Tris (pH 8.0), 137 mM NaCl, 1% glycerol, 0.1% SDS, 0.5% sodium deoxycholate, 1% Nonidet P-40, 2 mM EDTA, and protease inhibitor cocktail set III [Calbiochem, Darmstadt, Germany].) 20 min on ice. Lysates were clarified by centrifugation (20 min, 14800 rpm, 4°C). Samples (cell/viral lysate) were boiled at 95⁰C for 5 min with Roti load reducing loading buffer (Carl Roth, Karlsruhe, Germany) and subjected to SDS-PAGE followed by transfer (Semi-Dry Transfer Cell, Biorad, Munich, Germany) to a PVDF membrane (Merck Millipore, Schwalbach, Germany). Membranes were blocked with skimmed milk solution and probed with appropriate primary antibody, mouse anti-hemagglutinin (anti-HA) antibody (1:7,500 dilution, MMS-101P, Covance, Münster, Germany); mouse α-V5 antibody (1: 4000 dilution; Serotec); goat anti-GAPDH (C-terminus, 1:15,000 dilution, Everest Biotech, Oxfordshire, UK); mouse anti-α-tubulin antibody (1:4,000 dilution, clone B5-1-2; Sigma-Aldrich, Taufkirchen, Germany), mouse anti-capsid p24/p27 MAb AG3.0 [92] (1:250 dilution, NIH AIDS Reagents); rabbit anti S6 ribosomal protein (5G10; 1:10^3^ dilution in 5% BSA, Cell Signaling Technology, Leiden, The Netherlands). Secondary Abs.: anti-mouse (NA931V), anti-rabbit (NA934V) horseradish peroxidase (1:10^4^ dilution, GE Healthcare) and anti-goat IgG-HRP (1:10^4^ dilution, sc-2768, Santa Cruz Biotechnology, Heidelberg, Germany). Signals were visualized using ECL chemiluminescent reagent (GE Healthcare). To characterize the effect of the expression of A3 proteins and their mutants on LINE-1 (L1) reporter expression, HeLa-HA cells were lysed 48 h post-transfection using triple lysis buffer (20 mM Tris/HCl, pH 7.5; 150 mM NaCl; 10 mM EDTA; 0.1% SDS; 1% Triton X-100; 1% deoxycholate; 1x complete protease inhibitor cocktail [Roche]), clarified and 20 μg total protein were used for SDS-PAGE followed by electroblotting. HA-tagged A3 proteins and L1 ORF1p were detected using an anti-HA antibody (Cat.# MMS-101P; Covance Inc.) in a 1:5,000 dilution and the polyclonal rabbit-anti-L1 ORF1p antibody #984 [93] in a 1:2,000 dilution, respectively, in 1xPBS-T containing 5% milk powder. ß-actin expression (clone AC-74, 1:30,000 dilution, Sigma-Aldrich Chemie GmbH) served as a loading control.

### Differential DNA denaturation (3D) PCR

HEK293T cells were cultured in 6-well plates and infected with DNAse I (Thermo Fisher Scientific, Schwerte, Germany) treated viruses for 12 hours. Cells were harvested and washed in PBS, the total DNA was isolated using DNeasy DNA isolation kit (Qiagen, Hilden, Germany). A 714-bp fragment of the luciferase gene was amplified using the primers 5’-GATATGTGGATTTCGAGTCGTC-3’ and 5’-GTCATCGTCTTTCCGTGCTC-3’. For selective amplification of the hypermutated products, the PCR denaturation temperature was lowered stepwise from 87.6°C to 83.5°C (83.5°C, 84.2°C, 85.2°C, 86.3°C, 87.6°C) using a gradient thermocycler. The PCR parameters were as follows: (i) 95°C for 5 min; (ii) 40 cycles, with 1 cycle consisting of 83.5°C to 87.6°C for 30 s, 55°C for 30 s, 72°C for 1 min; (iii) 10 min at 72°C. PCRs were performed with Dream Taq DNA polymerase (Thermo Fisher Scientific). PCR products were stained with ethidium bromide. PCR product (smmA3C-like sample only) from the lowest denaturation temperature was cloned using CloneJET PCR Cloning Kit (Thermo Fisher Scientific) and sequenced. smmA3C-like protein-induced hypermutations of eleven independent clones were analysed with the Hypermut online tool (https://www.hiv.lanl.gov/content/sequence/HYPERMUT/hypermut.html) [94]. Mutated sequences (clones) carrying similar base changes were omitted and only the unique clones were presented for clarity.

### *In vitro* DNA cytidine deamination assay

A3 proteins expressed in transfected HEK293T cells or virion incorporated A3s used as input. Cell lysates were prepared with mild lysis buffer 48 h post plasmid transfection. Deamination reactions were performed as described [67, 95] in a 10 µL reaction volume containing 25 mM Tris pH 7.0, 2 µl of cell lysate and 100 fmol single-stranded DNA substrate (TTCA: 5’-GGATTGGTTGGTTATTTGTATAAGGAAGGTGGATTGAAGGTTCAAGAAGGTGATGGAAGTTATGTTTGGTAGATTGATGG). Samples were treated with 50 µg/ml RNAse A (Thermo Fisher Scientific). Reactions were incubated for 1 h at 37°C and the reaction was terminated by boiling at 95°C for 5 min. One fmol of the reaction mixture was used for PCR amplification Dream Taq polymerase (Thermo Fisher Scientific) 95°C for 3 min, followed by 30 cycles of 61°C for 30 s and 94°C for 30 s) using primers forward 5’-GGATTGGTTGGTTATTTGTATAAGGA and reverse 5’-CCATCAATCTACCAAACATAACTTCCA. PCR products were digested with MseI (NEB, Frankfurt/Main, Germany), and resolved on 15% PAGE, stained with ethidium bromide (7.5 μg/ml). As a positive control, substrate oligonucleotides with TTUA instead of TTCA were used to control the restriction enzyme digestion [65].

### L1 retrotransposition assay

Relative L1 retrotransposition activity was determined by applying a rapid dual-luciferase reporter based assay described previously [76]. Briefly, 2×10^5^ HeLa-HA cells were seeded per well of a six-well plate and transfected using Fugene-HD transfection reagent (Promega) according to the manufacturer’s protocol. Each well was cotransfected with 0.5 μg of the L1 retrotransposition reporter plasmid pYX017 or pYX015 [76] and 0.5 μg of pcDNA3.1 or wild-type or mutant A3 expression construct resuspended in 3 μl Fugene-HD transfection reagent and 100 μl GlutaMAX-I-supplemented Opti-MEM I reduced-serum medium (Thermo Fisher Scientific). Three days after transfection cultivation, the medium was replaced by complete DMEM containing 2.5 μg/ml puromycin, to select for the presence of the L1 reporter plasmid harboring a puroR-expression cassette. Next day, the medium was replaced once more by puromycin containing DMEM medium and 48 hours later, transfected cells were lysed to quantify dual-luciferase luminescence. Dual-luciferase luminescence measurement: Luminescence was measured using the Dual-Luciferase Reporter Assay System (Promega) following the manufacturer’s instructions. For assays in 6-well plates, 200 μl Passive Lysis Buffer was used to lyse cells in each well; for all assays, 20 μl lysate was transferred to a solid white 96-well plate, mixed with 50 μl Luciferase Assay Reagent II and firefly luciferase (Fluc) activity was quantified using the microplate luminometer Infinite 200PRO (Tecan, Männedorf, Switzerland). Renilla luciferase (Rluc) activity was subsequently read after mixing 50 μl Stop & Glo Reagent into the cell lysate containing Luciferase Assay Reagent II. Data were normalized as described in the results section. We routinely used the retrotransposition-defective L1RP/JM111 (located on pYX015) as the reference Fluc vector and set normalized luminescence ratio (NLR) resulting from cotransfection of pYX015 and pcDNA3.1(+) as 1.

### Protein sequence alignment and visualization

Sequence alignment of hA3C and smmA3C-like protein was done by using Clustal Omega (http://www.ebi.ac.uk/Tools/msa/clustalo/). The alignment file was then submitted to ESPript 3.0 [96] (espript.ibcp.fr) to calculate the similarity and identity of residues between both proteins and to build the alignment figure. Cartoon model of the crystal structure of A3C (PDB 3VOW) was constructed using PyMOL (PyMOL Molecular Graphics System version 1.5.0.4; Schrödinger, Portland, OR).

### Protein structural model building

The structural models of hA3C or C2 binding to ssDNA were generated by first aligning the X-ray crystal structure of rhA3G-NTD (PDB ID 5K82 [77]) onto the X-ray crystal structure of hA3F-CTD (PDB ID 5W2M [78]), the latter of which was co-crystallized with ssDNA. Subsequently, the hA3C X-ray crystal structure (PDB ID 3VOW [63]) was aligned onto the NTD of rhA3G, which is structurally similar to hA3C. The ssDNA and the interface region of hA3C were subsequently relaxed in the presence of each other using Maestro [97]. The same program was used to mutate hA3C to obtain the C2 and hA3C.S61P.S188I variants, which was again relaxed in the presence of the ssDNA. Similarly, we obtained a C2 ssDNA binding model based on the ssDNA-binding X-ray crystal structure of hA3A (PDB ID 5SWW [98]).

The alignment of the sequences of the crystal structures was generated with Probcons [99], accessed through Jalview [100], which was also used to visualize the alignment (Suppl. Fig. S9).

hA3C, C2, and hA3C.S61P.S188I were subjected to MD simulations. For this, the above-mentioned structures without the DNA were N- and C-terminally capped with ACE and NME, respectively. The three variants were protonated with PROPKA [101] according to pH 7.4, neutralized by adding counter ions, and solvated in an octahedral box of TIP3P water [102] with a minimal water shell of 12 Å around the solute. The Amber package of molecular simulation software [103] and the ff14SB force field [104] was used to perform the MD simulations. For the Zn^2+^-ions the Li-Merz parameters for two-fold positively charged metal ions [105] were used. To cope with long-range interactions, the “Particle Mesh Ewald” method [106] was used; the SHAKE algorithm [107] was applied to bonds involving hydrogen atoms. As hydrogen mass repartitioning [108] was utilized, the time step for all MD simulations was 4 fs with a direct-space, non-bonded cut-off of 8 Å. In the beginning, 17500 steps of steepest descent and conjugate gradient minimization were performed; during 2500, 10000, and 5000 steps positional harmonic restraints with force constants of 25 kcal mol^-1^ Å^-2^, 5 kcal mol^-1^ Å^-2^, and zero, respectively, were applied to the solute atoms. Thereafter, 50 ps of NVT (constant number of particles, volume, and temperature) MD simulations were conducted to heat up the system to 100 K, followed by 300 ps of NPT (constant number of particles, pressure, and temperature) MD simulations to adjust the density of the simulation box to a pressure of 1 atm and to heat the system to 300 K. During these steps, a harmonic potential with a force constant of 10 kcal mol^-1^ Å^-2^ was applied to the solute atoms. As the final step in thermalization, 300 ps of NVT-MD simulations were performed while gradually reducing the restraint forces on the solute atoms to zero within the first 100 ps of this step. Afterwards, five independent production runs of NVT-MD simulations with 2 μs length each were performed. For this, the starting temperatures of the MD simulations at the beginning of the thermalization were varied by a fraction of a Kelvin.

### Expression and purification of recombinant GST-tagged hA3C and hA3C.WE-RK from HEK293T cells

Recombinant C-terminal GST-tagged hA3C and hA3C.WE-RK were expressed in HEK293T cells and purified by affinity chromatography using Glutathione Sepharose 4B beads (GE Healthcare) as described previously [65]. Cells were lysed 48 h later with mild lysis buffer [50 mM Tris (pH 8), 1 mM PMSF, 10% glycerol, 0.8% NP-40, 150 mM NaCl, and 1X complete protease inhibitor and incubated with GST beads. After 2 h incubation at 4°C in end-over-end rotation, GST beads were washed twice with wash buffer containing 50 mM Tris (pH 8.0), 5 mM 2-ME, 10% glycerol and 500 mM NaCl. The bound GST hA3C and hA3C.WE-RK proteins were eluted with wash buffer containing 20 mM reduced glutathione. The proteins were 90-95% pure as checked on 15% SDS-PAGE followed by Coomassie blue staining. Protein concentrations were estimated by Bradford’s method.

### Electrophoretic mobility shift assay (EMSA) with hA3C-GST and hA3C.WE-RK-GST

EMSA was performed as described previously [65,79,109]. We mixed 20 fmol of 3′ biotinylated DNA (30-TTC-Bio-TEG purchased from Eurofins Genomics, Ebersberg Germany) with 10 mM Tris (pH − 7.5), 100 mM KCl, 10 mM MgCl2, 1 mM DTT, 2% glycerol, and the respective amount of recombinant proteins in a 15 μl reaction mixture, and incubated at room temperature for 30 min. The reaction mixture containing the protein–DNA complex were resolved on a 5% native PAGE gel on ice and transferred to a nylon membrane (Amersham Hybond-XL, GE healthcare) using 0.5 X TBE. After the transfer, the membrane containing protein–DNA complex were cross-linked by UV radiation with 312-nm bulb for 15 min. Chemiluminescent detection of biotinylated DNA was carried out according to the manufacturer’s instruction (Thermo Scientific LightShift Chemiluminescence EMSA Kit).

### Phylogenetic inference

In order to study the evolution of the A3-Z2 domains a representative set of 62 primate A3C, A3D, and A3F gene sequences were collected from GenBank (https://www.ncbi.nlm.nih.gov/genbank), as follows: 26 A3C sequences, 12 A3D sequences, and 21 A3F sequences. The phylogenetic relationships and divergence times among the species used were retrieved from http://www.timetree.org (Suppl. Fig. S10). A3 sequences from the northern tree shrew *Tupaia belangeri* were included as an outgroup to the primate ones. As A3D and A3F sequences contain each two Z2 domains, they were split into the corresponding N- and C-termini. The alignments were performed at the amino acid level using MAFFTv7.380 (http://mafft.cbrc.jp/alignment/software/) [110]. Phylogenetic inference was performed using RAxMLv8 [111], at either the nucleotide level under the GTR+Γ model or at the amino acid level under the LG+Γ model. Node support was evaluated applying 5,000 bootstrap cycles. Phylogenies at the nucleotide level were also calculated after introducing constraints in the tree, forcing monophyly of each clade A3D_N and C-termini, A3F_N and C-termini, New World monkeys A3C, and catarrhine A3C. Differences in maximum likelihood between alternative topologies for the same alignment were evaluated by the Shimodaira-Hasegawa test. Ancestral state reconstruction of amino acids in the loop A3_Z2 loop1 was performed only for the supported clades using RAxMLv8. A tanglegram with the two phylogenies was drawn with Dendroscope v3.6.3 [112]. Final layouts were done with Inkscape 0.92.4.

### Statistical analysis

Data were represented as the mean with SD in all bar diagrams. Statistically significant differences between two groups were analyzed using the unpaired Student’s t-test with GraphPad Prism version 5 (GraphPad Software, San Diego, CA, USA). A minimum *p-*value of 0.05 was considered as statistically significant.

## ACKNOWLEDGMENTS

We thank Wioletta Hörschken for excellent technical assistance. We thank Michael Emerman, Jens-Ove Heckel, Henning Hofmann, Yasumasa Iwatani, Nathanial R. Landau, Neeltje Kootstra, Bryan Cullen, Jonathan Stoye, Harald Wodrich, and Jörg Zielonka for reagents. The following reagents were obtained through the NIH AIDS Research and Reference Reagent Program, Division of AIDS, NIAID, NIH: a monoclonal antibody to HIV-1 p24 (AG3.0) from Jonathan Allan1. HG is grateful for computational support and infrastructure provided by the “Zentrum für Informations- und Medientechnologie” (ZIM) at the Heinrich-Heine-University Düsseldorf and the computing time provided by the John von Neumann Institute for Computing (NIC) to HG on the supercomputer JUWELS at Jülich Supercomputing Centre (JSC) (user ID: HKF7).

## AUTHOR CONTRIBUTIONS

Conceptualization: AAJV and CM

Data curation: AAJV, KB, CGWG, FB, UH, CK, SB, GGS, IGB, DH, and HG

Formal analysis: AAJV, KB, CGWG, FB, ZZ, AS, UH, SB, CK, GGS, IGB, DH, HG, and CM

Funding acquisition: AAJV, CM, and HG

Investigation: AAJV, KB, CGWG, FB, UH, CK, HG, and IGB

Methodology: AAJV, KB, CGWG, SB, HG, and CM

Project administration: CM

Resources: GGS, IGB, HG, and CM

Supervision: CM, DH, and AAJV

Validation: AAJV and CM

Visualization: AAJV, KB, CGWG, FB, CK, GGS, IGB, HG, and CM

Writing – original draft: AAJV

Writing – review and editing: AAJV, KB, CGWG, FB, ZZ, AS, UH, CK, SB, GGS, DH, IGB, HG, and CM

## SUPPLEMENTARY FIGURES

**S1 Fig.**
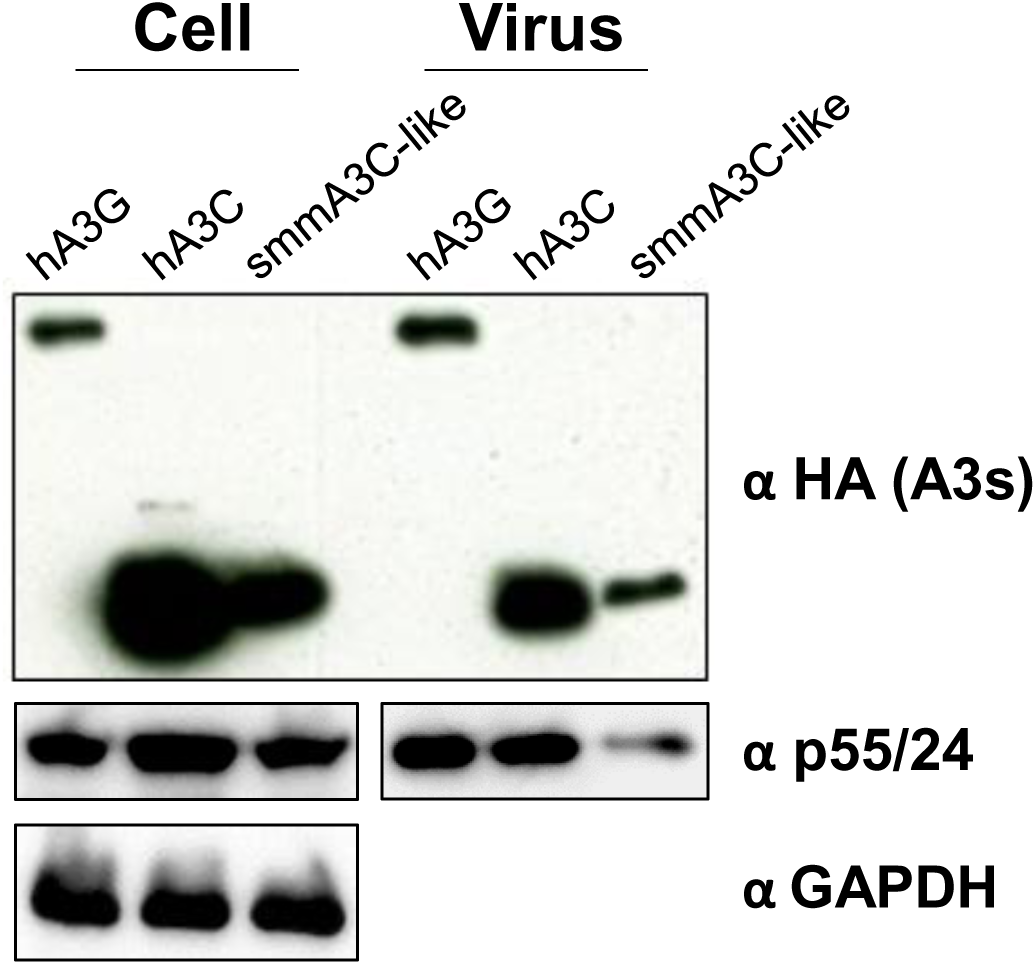
Viral incorporation of A3s. Immunoblot analysis of HA-tagged hA3G, hA3C, and smmA3C-like protein in lysates of transfected cells (Cell) and HIV-1Δ*vif* viral particles (Virus). A3s were detected using anti-HA and anti p24 antibodies, respectively. GAPDH served as a loading control. “α” represents anti.

**S2 Fig.**
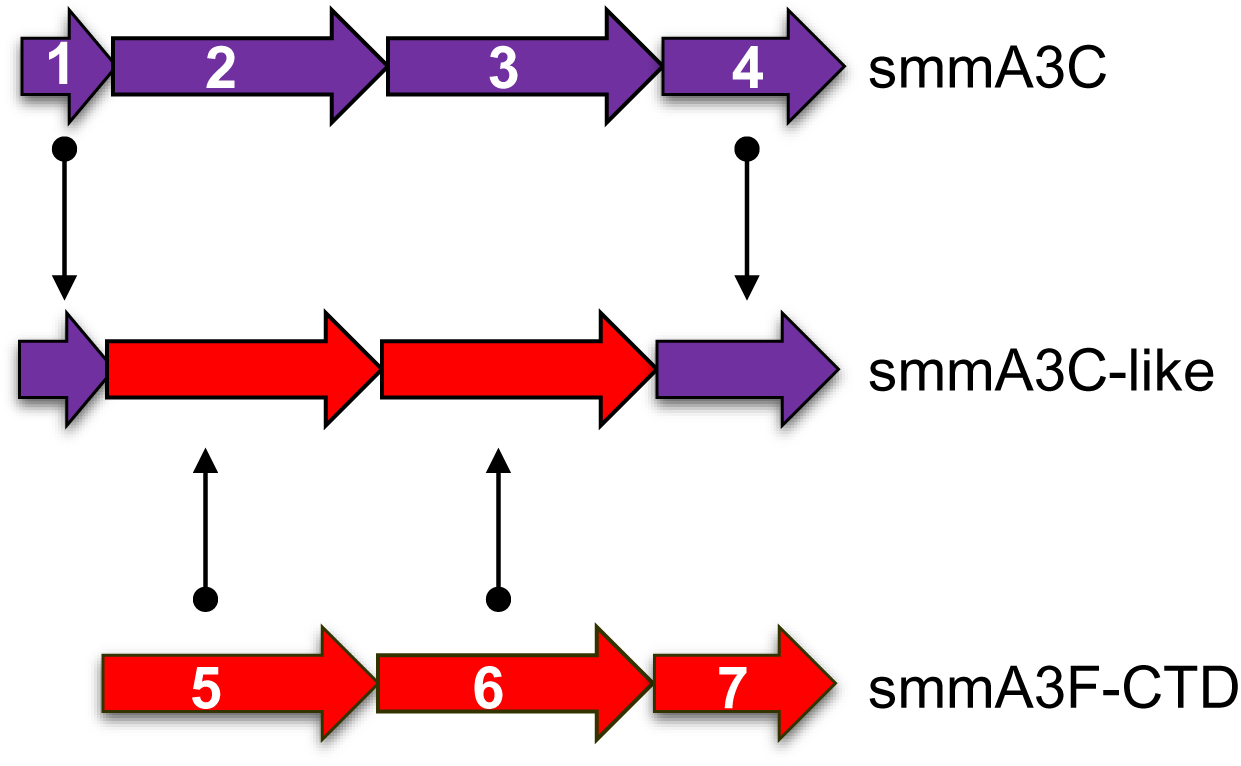
The ORF of smmA3C-like is a hybrid sequence made by exonic sequences of smmA3C and smmA3F. Arrangement of exons of the ORFs that produce smmA3C-like protein, smmA3C, and smmA3F-CTD. Purple and red colors indicate exons derived from smmA3C and smmA3F, respectively. Note that smmA3F-CTD is encoded by exons 5, 6, and 7 of smmA3F.

**S3 Fig.**
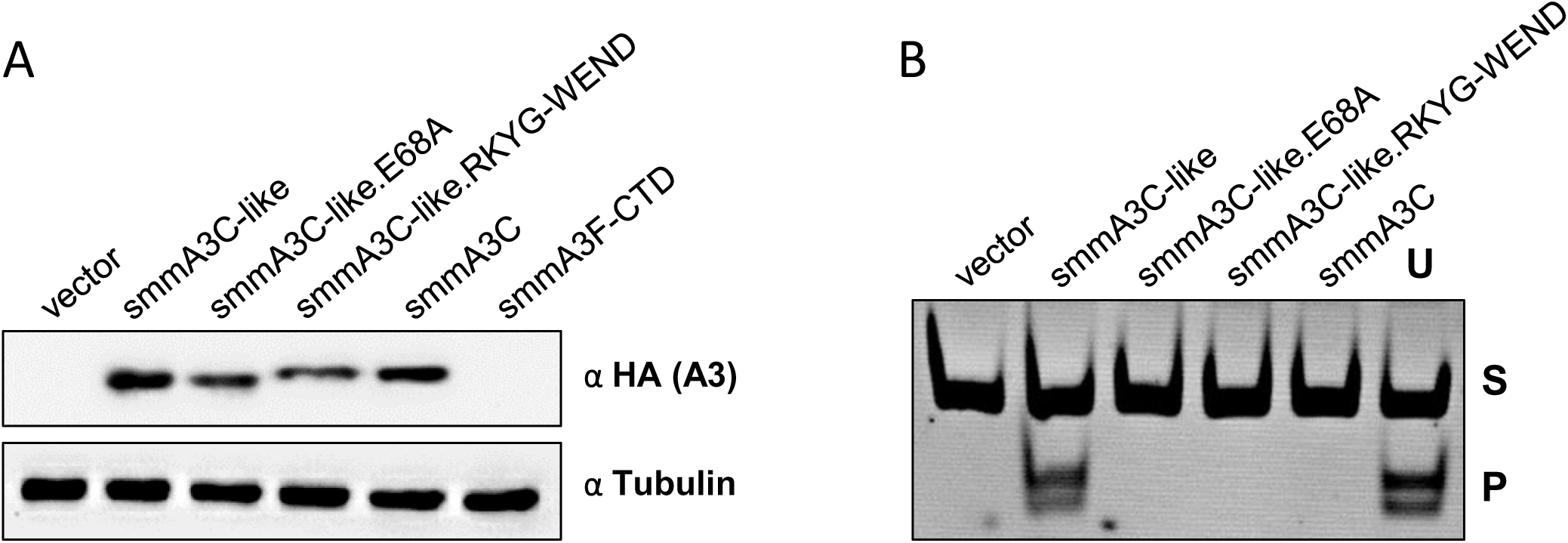
Expression and deamination activity of smmA3C and smmA3F-CTD. (A) HEK293T cells were transfected with expression plasmids encoding smmA3C, smmA3F-CTD, smmA3C-like protein and their mutants. Immunoblot stained with anti-HA antibody, shows the amount of A3s in cell lysates. Tubulin served as a loading control. “α” represent anti. (B) To examine the catalytic activity of smmA3C, smmA3C-like protein, and their variants, *in vitro* deamination assay was performed using lysates of cells that were previously transfected with respective expression plasmids. RNAse A-treatment was included; oligonucleotide containing uracil (U) instead of cytosine served as a marker to denote the migration of deaminated product after restriction enzyme cleavage. S-substrate, P-product.

**S4 Fig.**
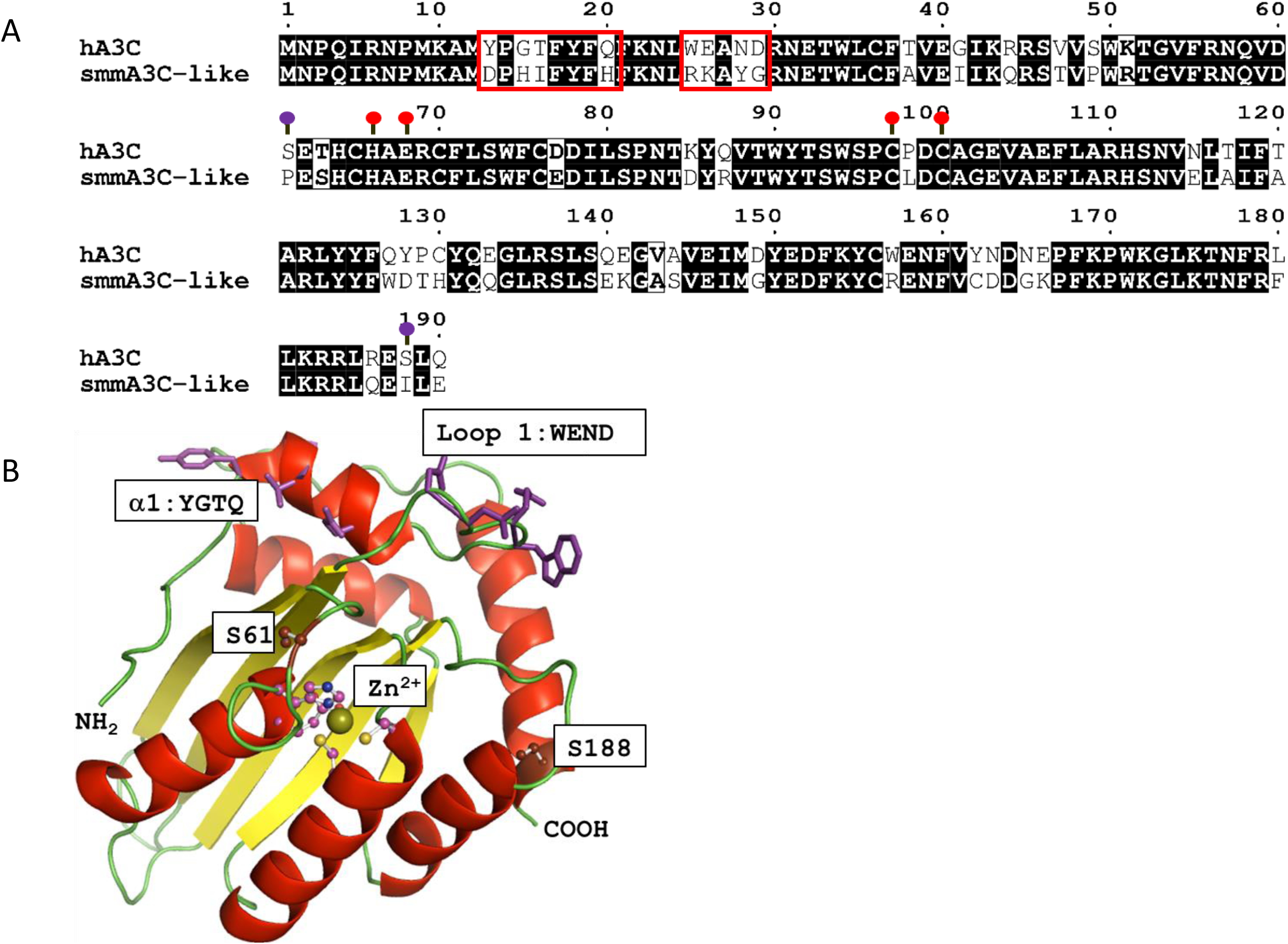
Protein sequence and structure information of hA3C and smmA3C-like protein. (A) Sequence alignment of hA3C and smmA3C-like protein was done by using Clustal Omega and ESPript 3.0. Motif 1 (YGTQ) and motif 2 (WEND) are marked with red boxes, red lollipops indicate active site amino acids H66, E68, C97 and C100, while S61 and S188 are colored in purple. (B) Ribbon model of the crystal structure of A3C (PDB 3VOW) depicting the spatial arrangements of helix α1 (YGTQ motif) and loop 1 (WEND motif). Residues of both motifs are presented in purple. Key residues S61, S188, and zinc-coordinating active site residues are denoted as ball and sticks. Sphere represents Zn^2+^ ion.

**S5 Fig.**
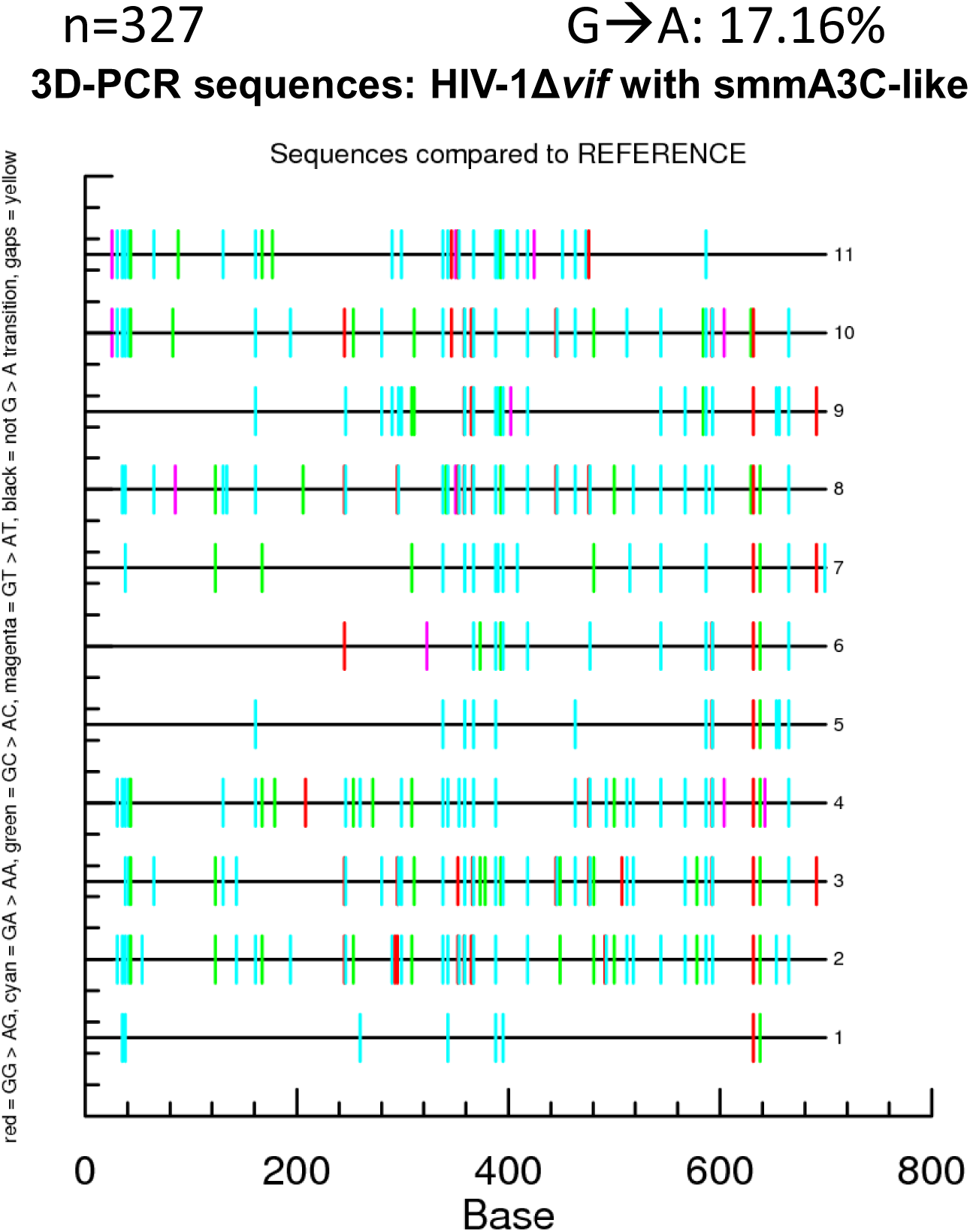
smmA3C-like protein mediated hypermutation of HIV-1Δ*vif* DNA. To characterize and quantify mutations induced by smmA3C-like protein, the PCR product obtained at the lowest denaturing temperature (shown in Fig. 1C and marked with a red asterisk (*)) was cloned and a number of independent clones were sequenced and analyzed. The mutation pattern is presented using hypermut tool (note: identical clones were omitted). Each of the sequences is given as a horizontal line. Mutations were denoted as small vertical lines. Red, cyan, magenta, green and black colored lines represent GG-to-AG, GA-to-AA, GT-to-AT, GC-to-AC, and non-G-to-A mutations, respectively. The overall mutation load (G→A) was given in the top ride side as %. ‘n’ denotes the total number of independent G→A mutations found in independent sequences.

**S6 Fig.**
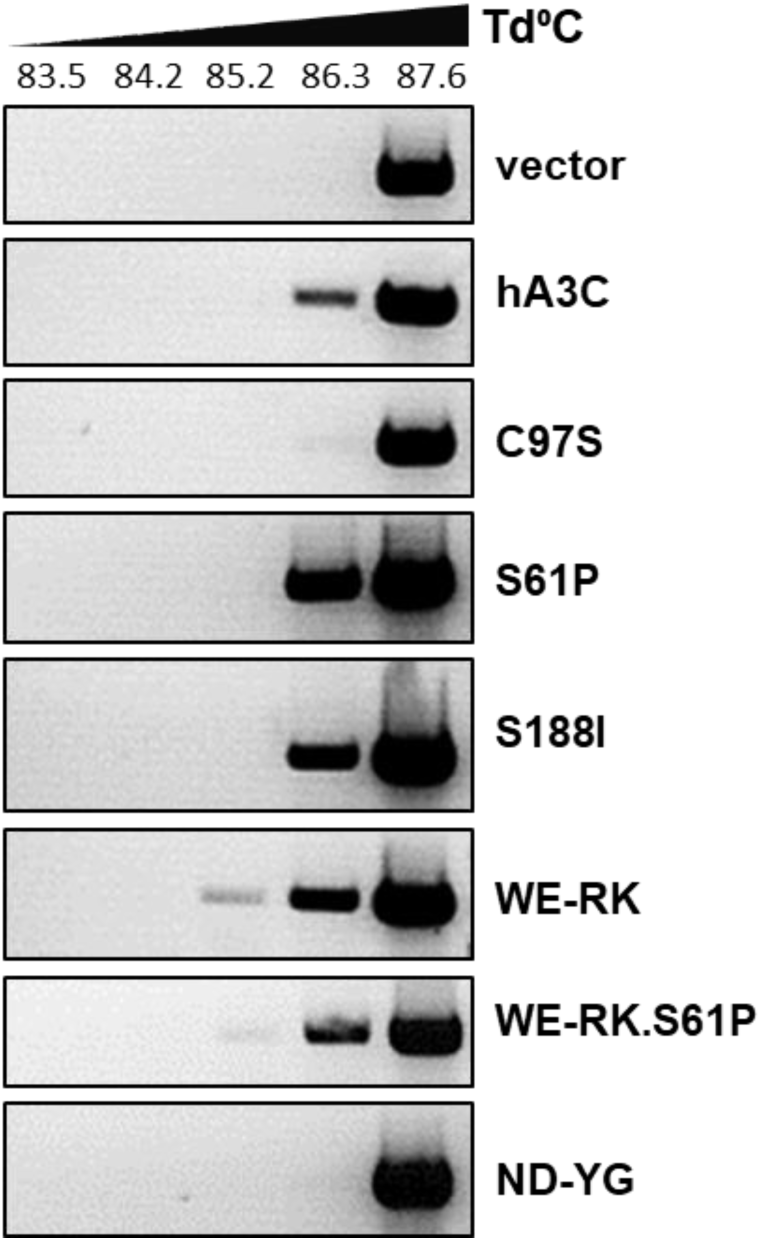
3D-PCR analysis of hA3C mutants. HIV-1Δ*vif* produced together with hA3C, its variants (C97S, S61P, S188I, WE-RK, WE-RK.S61P, ND-YG), or vector controls were used to transduce HEK293T cells. Total DNA was extracted and a 714-bp fragment of reporter viral DNA was selectively amplified using 3D-PCR. Td = denaturation.

**S7 Fig.**
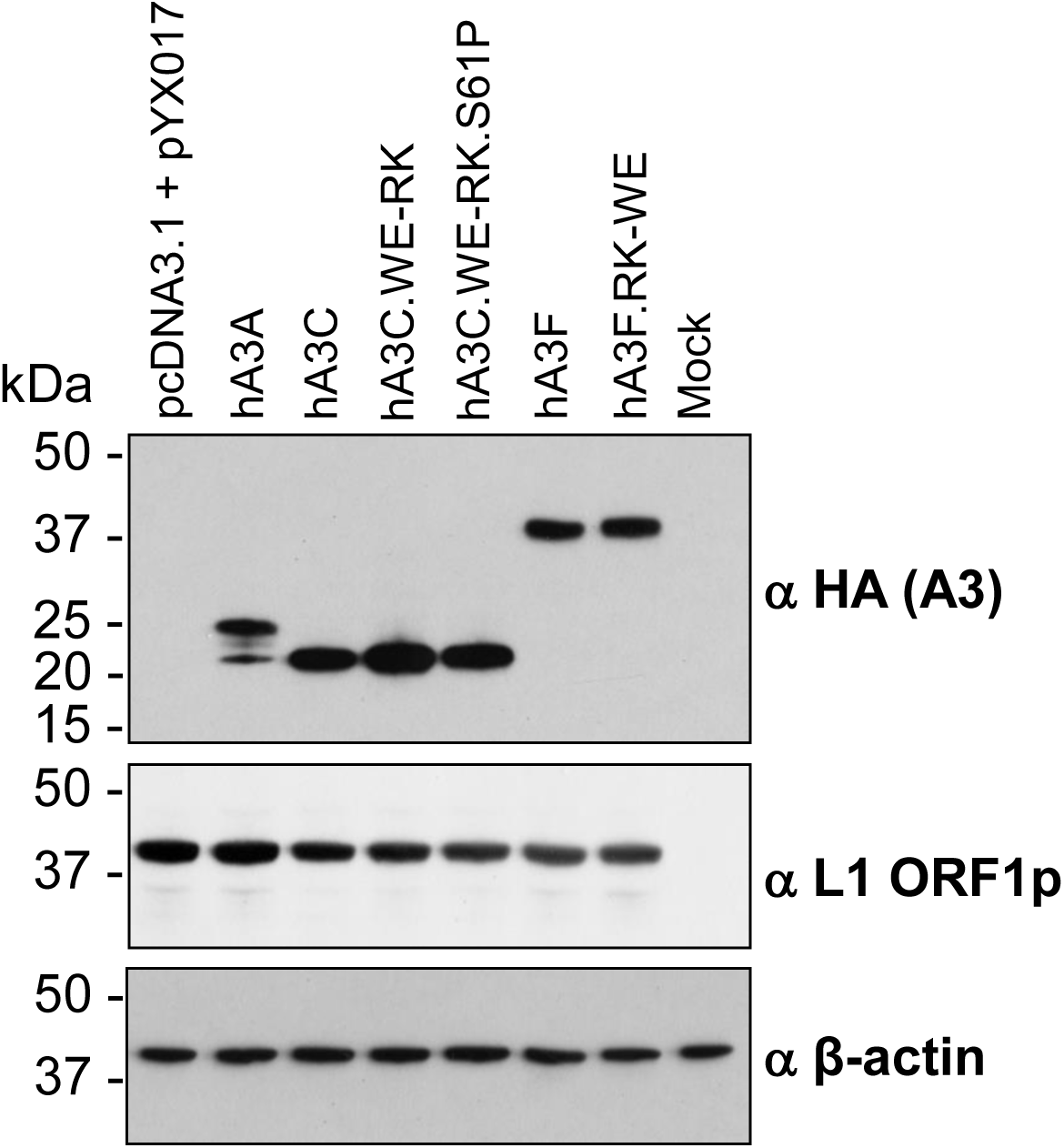
Expression of transfected A3- and L1-plasmids in HeLa cells. Immunoblot analysis of the expression of A3C and its mutants WE-RK, WE-RK.S61P, A3A, A3F, and its mutant RK-WE after co-transfection with the L1 reporter pYX017 into HeLa-HA cells. Expression of the L1 reporter and A3 proteins was analyzed using antibodies against L1 ORF1p (α-ORF1p) or α-HA, respectively. β-actin protein levels were analyzed as loading controls. Cell extracts from mock-transfected HeLa-HA cells served as negative controls for L1-ORF1p expression and HA-tagged A3 expression.

**S8 Fig.**
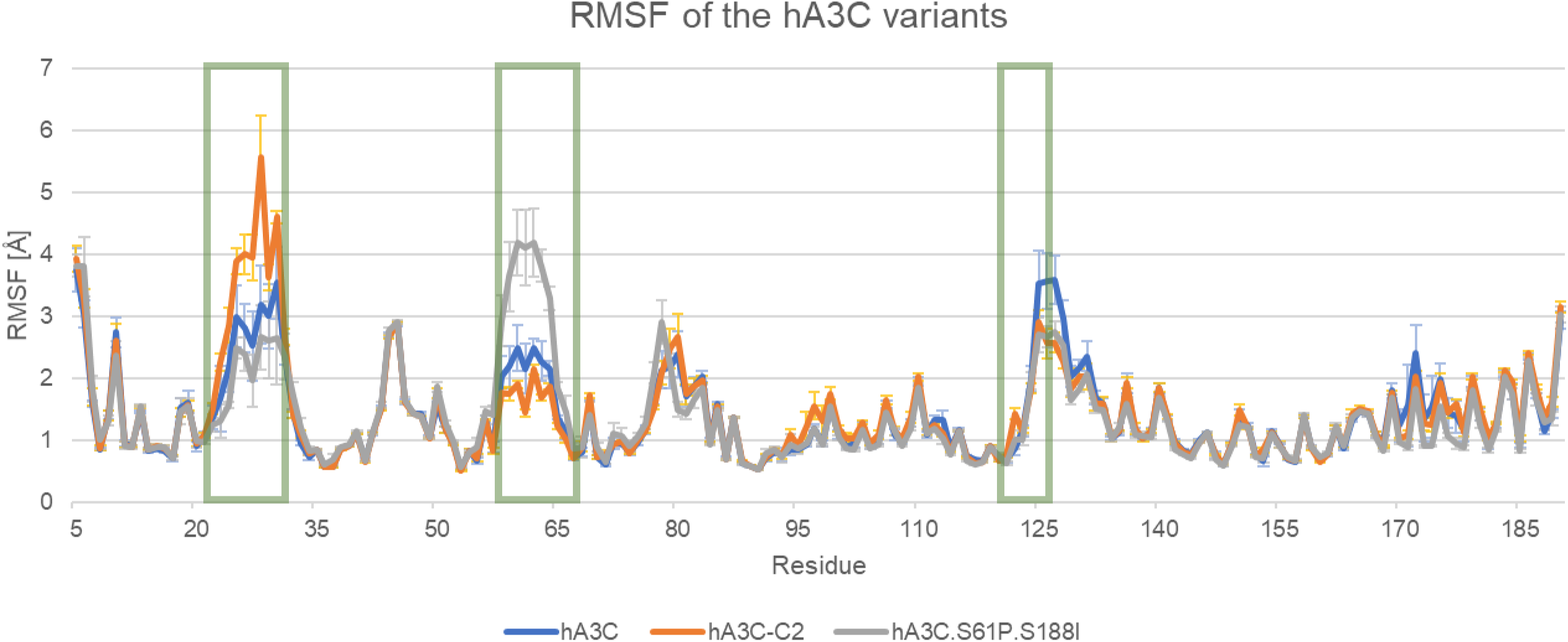
Root mean square fluctuations of the residues in the hA3C, C2, and hA3C.S61P.S188I variants over the course of 2 µs MD simulations. The RMSF are given as means over five independent replicas for each variant ± SEM for all residues present in the X-ray structure of A3C (PDB ID 3VOW). The residues lining the active center are marked by green boxes.

**S9 Fig.**
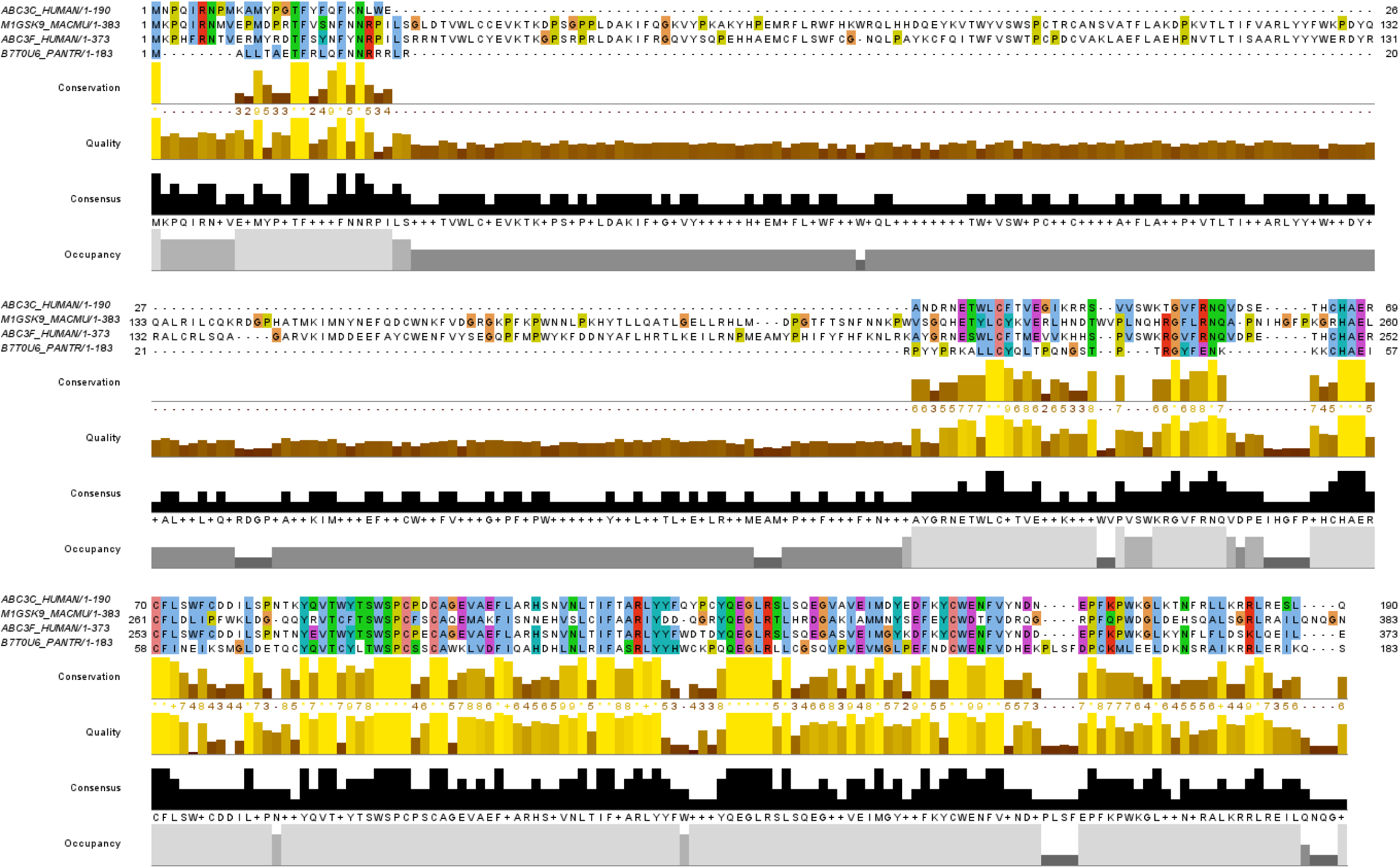
Multiple sequence alignment of human A3C (ABC3C_HUMAN), rhesus macaque A3G (M1GSK9_MACMU), human A3F (ABC3F_HUMAN), and chimpanzee A3H (B7T0U6_PANTR), in order. The coloring of the residues depends on the physicochemical properties of conserved residues.

**S10 Fig.**
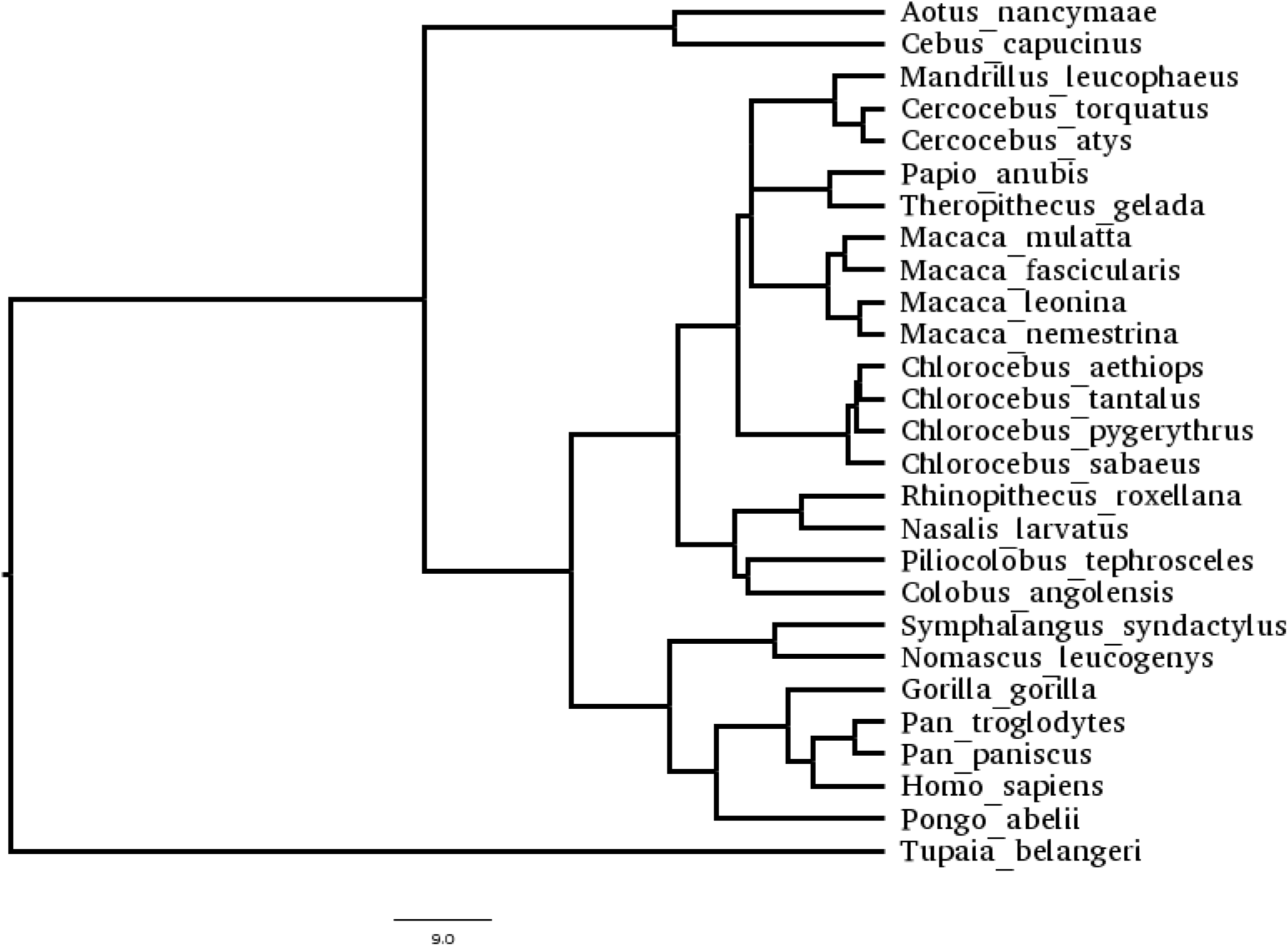
Phylogenetic relationships and divergence times among the species used to collect the A3Z2 sequences, were retrieved from http://www.timetree.org.

**S1 table:**
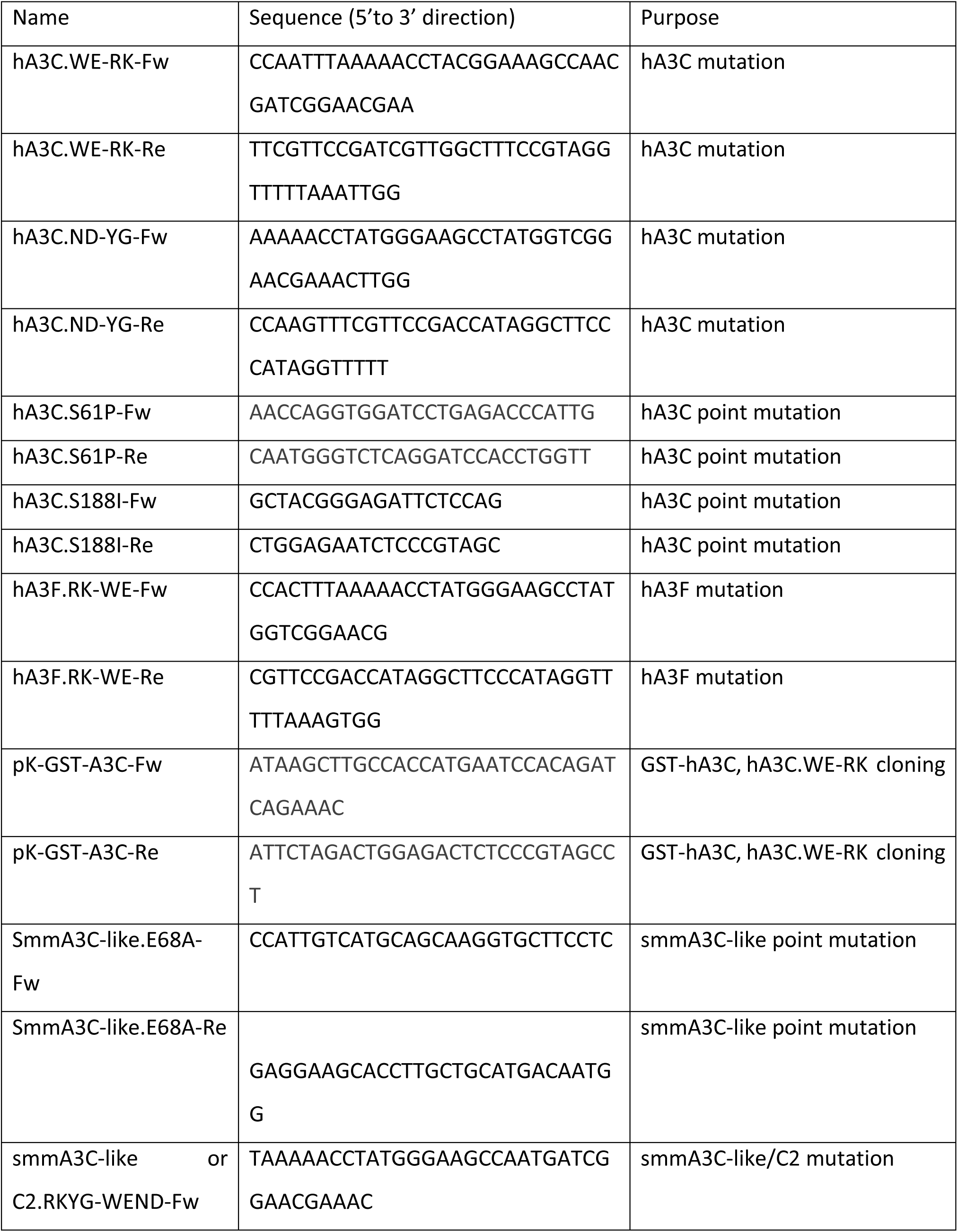
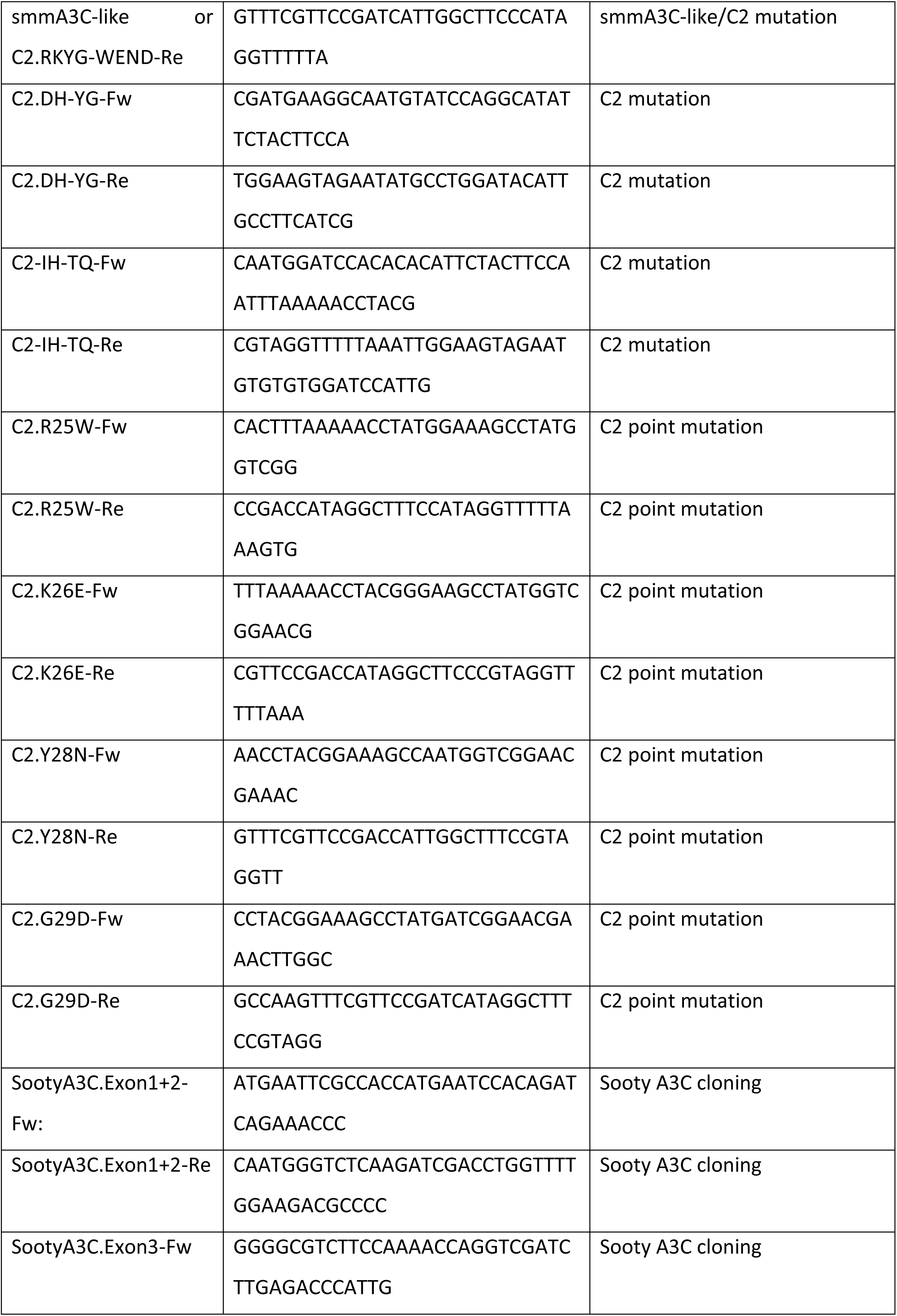
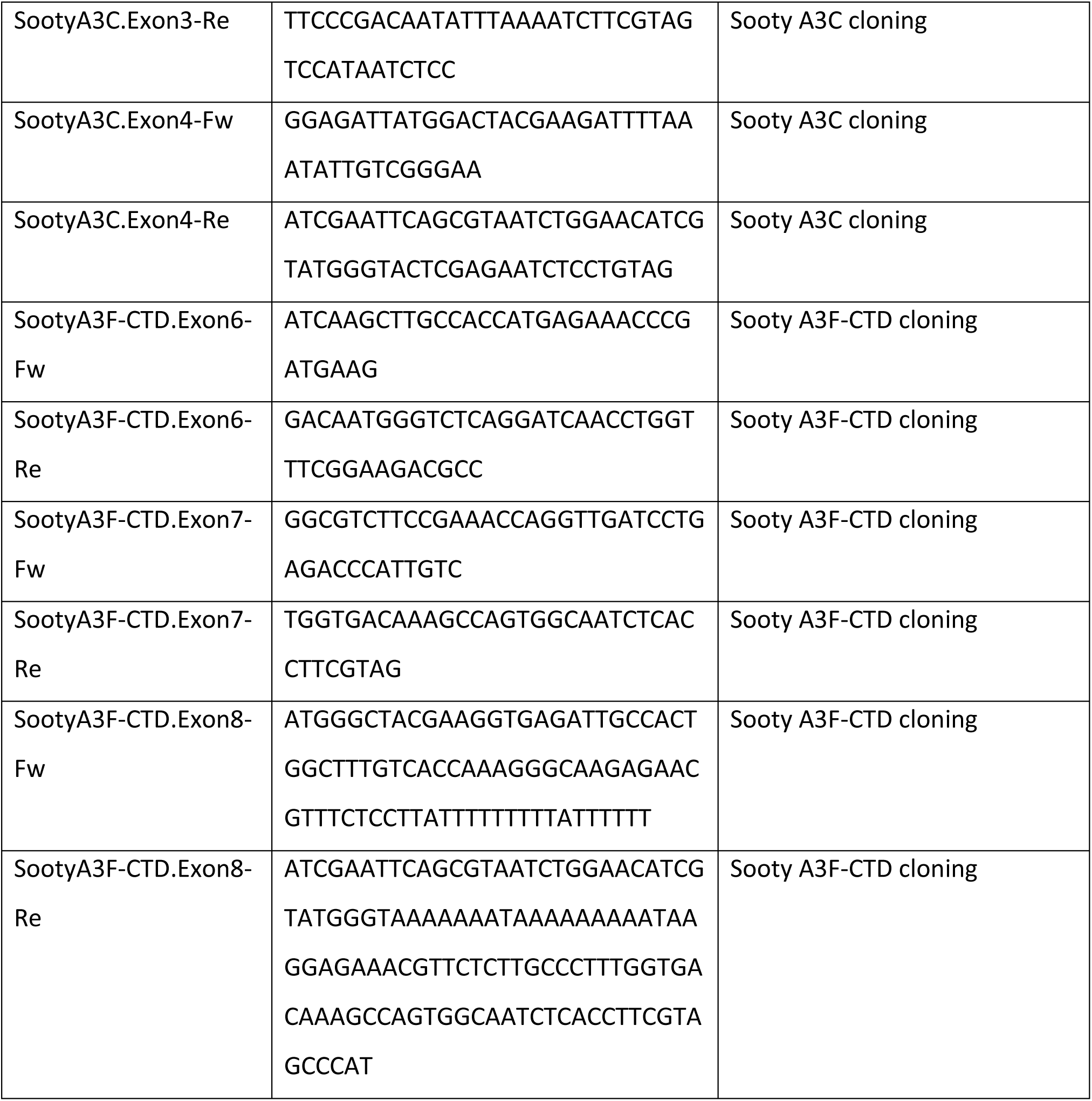
List of oligonucleotides used in the study.

## REFERENCES

1. Goila-Gaur R, Strebel K (2008) HIV-1 Vif, APOBEC, and intrinsic immunity. Retrovirology 5: 51.

2. Harris RS, Dudley JP (2015) APOBECs and virus restriction. Virology 479-480: 131–145.

3. Salter JD, Bennett RP, Smith HC (2016) The APOBEC Protein Family: United by Structure, Divergent in Function. Trends Biochem Sci 41: 578–594.

4. Silvas TV, Schiffer CA (2019) APOBEC3s: DNA-editing human cytidine deaminases. Protein Sci 28: 1552–1566.

5. Jarmuz A, Chester A, Bayliss J, Gisbourne J, Dunham I, et al. (2002) An anthropoid-specific locus of orphan C to U RNA-editing enzymes on chromosome 22. Genomics 79: 285–296.

6. Münk C, Willemsen A, Bravo IG (2012) An ancient history of gene duplications, fusions and losses in the evolution of APOBEC3 mutators in mammals. BMC Evol Biol 12: 71.

7. LaRue RS, Jonsson SR, Silverstein KA, Lajoie M, Bertrand D, et al. (2008) The artiodactyl APOBEC3 innate immune repertoire shows evidence for a multi-functional domain organization that existed in the ancestor of placental mammals. BMC Mol Biol 9: 104.

8. Sheehy AM, Gaddis NC, Choi JD, Malim MH (2002) Isolation of a human gene that inhibits HIV-1 infection and is suppressed by the viral Vif protein. Nature 418: 646–650.

9. Zhang H, Yang B, Pomerantz RJ, Zhang C, Arunachalam SC, et al. (2003) The cytidine deaminase CEM15 induces hypermutation in newly synthesized HIV-1 DNA. Nature 424: 94–98.

10. Bishop KN, Holmes RK, Sheehy AM, Davidson NO, Cho SJ, et al. (2004) Cytidine deamination of retroviral DNA by diverse APOBEC proteins. Curr Biol 14: 1392–1396.

11. Vasudevan AA, Smits SH, Hoppner A, Häussinger D, Koenig BW, et al. (2013) Structural features of antiviral DNA cytidine deaminases. Biol Chem 394: 1357–1370.

12. Zennou V, Perez-Caballero D, Gottlinger H, Bieniasz PD (2004) APOBEC3G incorporation into human immunodeficiency virus type 1 particles. J Virol 78: 12058–12061.

13. Luo K, Liu B, Xiao Z, Yu Y, Yu X, et al. (2004) Amino-terminal region of the human immunodeficiency virus type 1 nucleocapsid is required for human APOBEC3G packaging. J Virol 78: 11841–11852.

14. Svarovskaia ES, Xu H, Mbisa JL, Barr R, Gorelick RJ, et al. (2004) Human apolipoprotein B mRNA-editing enzyme-catalytic polypeptide-like 3G (APOBEC3G) is incorporated into HIV-1 virions through interactions with viral and nonviral RNAs. J Biol Chem 279: 35822–35828.

15. Huthoff H, Malim MH (2007) Identification of amino acid residues in APOBEC3G required for regulation by human immunodeficiency virus type 1 Vif and Virion encapsidation. J Virol 81: 3807–3815.

16. Schafer A, Bogerd HP, Cullen BR (2004) Specific packaging of APOBEC3G into HIV-1 virions is mediated by the nucleocapsid domain of the gag polyprotein precursor. Virology 328: 163–168.

17. Burnett A, Spearman P (2007) APOBEC3G multimers are recruited to the plasma membrane for packaging into human immunodeficiency virus type 1 virus-like particles in an RNA-dependent process requiring the NC basic linker. J Virol 81: 5000–5013.

18. Browne EP, Allers C, Landau NR (2009) Restriction of HIV-1 by APOBEC3G is cytidine deaminase-dependent. Virology 387: 313–321.

19. Harris RS, Bishop KN, Sheehy AM, Craig HM, Petersen-Mahrt SK, et al. (2003) DNA deamination mediates innate immunity to retroviral infection. Cell 113: 803–809.

20. Yu Q, König R, Pillai S, Chiles K, Kearney M, et al. (2004) Single-strand specificity of APOBEC3G accounts for minus-strand deamination of the HIV genome. Nat Struct Mol Biol 11: 435–442.

21. Mangeat B, Turelli P, Caron G, Friedli M, Perrin L, et al. (2003) Broad antiretroviral defence by human APOBEC3G through lethal editing of nascent reverse transcripts. Nature 424: 99–103.

22. Iwatani Y, Chan DS, Wang F, Maynard KS, Sugiura W, et al. (2007) Deaminase-independent inhibition of HIV-1 reverse transcription by APOBEC3G. Nucleic Acids Res 35: 7096–7108.

23. Holmes RK, Koning FA, Bishop KN, Malim MH (2007) APOBEC3F can inhibit the accumulation of HIV-1 reverse transcription products in the absence of hypermutation. Comparisons with APOBEC3G. J Biol Chem 282: 2587–2595.

24. Münk C, Jensen BE, Zielonka J, Häussinger D, Kamp C (2012) Running Loose or Getting Lost: How HIV-1 Counters and Capitalizes on APOBEC3-Induced Mutagenesis through Its Vif Protein. Viruses 4: 3132–3161.

25. Bishop KN, Holmes RK, Malim MH (2006) Antiviral potency of APOBEC proteins does not correlate with cytidine deamination. J Virol 80: 8450–8458.

26. Mbisa JL, Bu W, Pathak VK (2010) APOBEC3F and APOBEC3G inhibit HIV-1 DNA integration by different mechanisms. J Virol 84: 5250–5259.

27. Strebel K (2005) APOBEC3G & HTLV-1: inhibition without deamination. Retrovirology 2: 37.

28. Mehle A, Strack B, Ancuta P, Zhang C, McPike M, et al. (2004) Vif overcomes the innate antiviral activity of APOBEC3G by promoting its degradation in the ubiquitin-proteasome pathway. J Biol Chem 279: 7792–7798.

29. Sheehy AM, Gaddis NC, Malim MH (2003) The antiretroviral enzyme APOBEC3G is degraded by the proteasome in response to HIV-1 Vif. Nat Med 9: 1404–1407.

30. Yu X, Yu Y, Liu B, Luo K, Kong W, et al. (2003) Induction of APOBEC3G ubiquitination and degradation by an HIV-1 Vif-Cul5-SCF complex. Science 302: 1056–1060.

31. Mariani R, Chen D, Schrofelbauer B, Navarro F, König R, et al. (2003) Species-specific exclusion of APOBEC3G from HIV-1 virions by Vif. Cell 114: 21–31.

32. Bogerd HP, Doehle BP, Wiegand HL, Cullen BR (2004) A single amino acid difference in the host APOBEC3G protein controls the primate species specificity of HIV type 1 virion infectivity factor. Proc Natl Acad Sci U S A 101: 3770–3774.

33. Mangeat B, Turelli P, Liao S, Trono D (2004) A single amino acid determinant governs the species-specific sensitivity of APOBEC3G to Vif action. J Biol Chem 279: 14481–14483.

34. Zhang W, Huang M, Wang T, Tan L, Tian C, et al. (2008) Conserved and non-conserved features of HIV-1 and SIVagm Vif mediated suppression of APOBEC3 cytidine deaminases. Cell Microbiol 10: 1662–1675.

35. Smith JL, Pathak VK (2010) Identification of specific determinants of human APOBEC3F, APOBEC3C, and APOBEC3DE and African green monkey APOBEC3F that interact with HIV-1 Vif. J Virol 84: 12599–12608.

36. Dang Y, Wang X, Esselman WJ, Zheng YH (2006) Identification of APOBEC3DE as another antiretroviral factor from the human APOBEC family. J Virol 80: 10522–10533.

37. Wiegand HL, Doehle BP, Bogerd HP, Cullen BR (2004) A second human antiretroviral factor, APOBEC3F, is suppressed by the HIV-1 and HIV-2 Vif proteins. EMBO J 23: 2451–2458.

38. Zheng YH, Irwin D, Kurosu T, Tokunaga K, Sata T, et al. (2004) Human APOBEC3F is another host factor that blocks human immunodeficiency virus type 1 replication. J Virol 78: 6073–6076.

39. Hultquist JF, Lengyel JA, Refsland EW, LaRue RS, Lackey L, et al. (2011) Human and rhesus APOBEC3D, APOBEC3F, APOBEC3G, and APOBEC3H demonstrate a conserved capacity to restrict Vif-deficient HIV-1. J Virol 85: 11220–11234.

40. Burns MB, Lackey L, Carpenter MA, Rathore A, Land AM, et al. (2013) APOBEC3B is an enzymatic source of mutation in breast cancer. Nature 494: 366–370.

41. Roberts SA, Lawrence MS, Klimczak LJ, Grimm SA, Fargo D, et al. (2013) An APOBEC cytidine deaminase mutagenesis pattern is widespread in human cancers. Nat Genet 45: 970–976.

42. Henderson S, Fenton T (2015) APOBEC3 genes: retroviral restriction factors to cancer drivers. Trends Mol Med 21: 274–284.

43. Swanton C, McGranahan N, Starrett GJ, Harris RS (2015) APOBEC Enzymes: Mutagenic Fuel for Cancer Evolution and Heterogeneity. Cancer Discov 5: 704–712.

44. Green AM, Weitzman MD (2019) The spectrum of APOBEC3 activity: From anti-viral agents to anti-cancer opportunities. DNA Repair (Amst) 83: 102700.

45. Olson ME, Harris RS, Harki DA (2018) APOBEC Enzymes as Targets for Virus and Cancer Therapy. Cell Chem Biol 25: 36–49.

46. Muckenfuss H, Hamdorf M, Held U, Perkovic M, Lower J, et al. (2006) APOBEC3 proteins inhibit human LINE-1 retrotransposition. J Biol Chem 281: 22161–22172.

47. Yu Q, Chen D, König R, Mariani R, Unutmaz D, et al. (2004) APOBEC3B and APOBEC3C are potent inhibitors of simian immunodeficiency virus replication. J Biol Chem 279: 53379–53386.

48. Langlois MA, Beale RC, Conticello SG, Neuberger MS (2005) Mutational comparison of the single-domained APOBEC3C and double-domained APOBEC3F/G anti-retroviral cytidine deaminases provides insight into their DNA target site specificities. Nucleic Acids Res 33: 1913–1923.

49. Suspene R, Guetard D, Henry M, Sommer P, Wain-Hobson S, et al. (2005) Extensive editing of both hepatitis B virus DNA strands by APOBEC3 cytidine deaminases in vitro and in vivo. Proc Natl Acad Sci U S A 102: 8321–8326.

50. Baumert TF, Rosler C, Malim MH, von Weizsacker F (2007) Hepatitis B virus DNA is subject to extensive editing by the human deaminase APOBEC3C. Hepatology 46: 682–689.

51. Vartanian JP, Guetard D, Henry M, Wain-Hobson S (2008) Evidence for editing of human papillomavirus DNA by APOBEC3 in benign and precancerous lesions. Science 320: 230–233.

52. Stauch B, Hofmann H, Perkovic M, Weisel M, Kopietz F, et al. (2009) Model structure of APOBEC3C reveals a binding pocket modulating ribonucleic acid interaction required for encapsidation. Proc Natl Acad Sci U S A 106: 12079–12084.

53. Ahasan MM, Wakae K, Wang Z, Kitamura K, Liu G, et al. (2015) APOBEC3A and 3C decrease human papillomavirus 16 pseudovirion infectivity. Biochem Biophys Res Commun 457: 295–299.

54. Suspene R, Aynaud MM, Koch S, Pasdeloup D, Labetoulle M, et al. (2011) Genetic editing of herpes simplex virus 1 and Epstein-Barr herpesvirus genomes by human APOBEC3 cytidine deaminases in culture and in vivo. J Virol 85: 7594–7602.

55. Perkovic M, Schmidt S, Marino D, Russell RA, Stauch B, et al. (2009) Species-specific inhibition of APOBEC3C by the prototype foamy virus protein bet. J Biol Chem 284: 5819–5826.

56. Horn AV, Klawitter S, Held U, Berger A, Vasudevan AA, et al. (2014) Human LINE-1 restriction by APOBEC3C is deaminase independent and mediated by an ORF1p interaction that affects LINE reverse transcriptase activity. Nucleic Acids Res 42: 396–416.

57. Hultquist JF, Binka M, LaRue RS, Simon V, Harris RS (2012) Vif proteins of human and simian immunodeficiency viruses require cellular CBFbeta to degrade APOBEC3 restriction factors. J Virol 86: 2874–2877.

58. Bonvin M, Achermann F, Greeve I, Stroka D, Keogh A, et al. (2006) Interferon-inducible expression of APOBEC3 editing enzymes in human hepatocytes and inhibition of hepatitis B virus replication. Hepatology 43: 1364–1374.

59. Refsland EW, Hultquist JF, Harris RS (2012) Endogenous origins of HIV-1 G-to-A hypermutation and restriction in the nonpermissive T cell line CEM2n. PLoS Pathog 8: e1002800.

60. Bourara K, Liegler TJ, Grant RM (2007) Target cell APOBEC3C can induce limited G-to-A mutation in HIV-1. PLoS Pathog 3: 1477–1485.

61. Refsland EW, Stenglein MD, Shindo K, Albin JS, Brown WL, et al. (2010) Quantitative profiling of the full APOBEC3 mRNA repertoire in lymphocytes and tissues: implications for HIV-1 restriction. Nucleic Acids Res 38: 4274–4284.

62. Abdel-Mohsen M, Raposo RA, Deng X, Li M, Liegler T, et al. (2013) Expression profile of host restriction factors in HIV-1 elite controllers. Retrovirology 10: 106.

63. Kitamura S, Ode H, Nakashima M, Imahashi M, Naganawa Y, et al. (2012) The APOBEC3C crystal structure and the interface for HIV-1 Vif binding. Nat Struct Mol Biol 19: 1005–1010.

64. Zhang Z, Gu Q, Jaguva Vasudevan AA, Jeyaraj M, Schmidt S, et al. (2016) Vif Proteins from Diverse Human Immunodeficiency Virus/Simian Immunodeficiency Virus Lineages Have Distinct Binding Sites in A3C. J Virol 90: 10193–10208.

65. Jaguva Vasudevan AA, Hofmann H, Willbold D, Häussinger D, Koenig BW, et al. (2017) Enhancing the Catalytic Deamination Activity of APOBEC3C Is Insufficient to Inhibit Vif-Deficient HIV-1. J Mol Biol 429: 1171–1191.

66. Suspene R, Henry M, Guillot S, Wain-Hobson S, Vartanian JP (2005) Recovery of APOBEC3-edited human immunodeficiency virus G->A hypermutants by differential DNA denaturation PCR. J Gen Virol 86: 125–129.

67. Nowarski R, Britan-Rosich E, Shiloach T, Kotler M (2008) Hypermutation by intersegmental transfer of APOBEC3G cytidine deaminase. Nat Struct Mol Biol 15: 1059–1066.

68. Jaguva Vasudevan AA, Kreimer U, Schulz WA, Krikoni A, Schumann GG, et al. (2018) APOBEC3B Activity Is Prevalent in Urothelial Carcinoma Cells and Only Slightly Affected by LINE-1 Expression. Front Microbiol 9: 2088.

69. Wittkopp CJ, Adolph MB, Wu LI, Chelico L, Emerman M (2016) A Single Nucleotide Polymorphism in Human APOBEC3C Enhances Restriction of Lentiviruses. PLoS Pathog 12: e1005865.

70. Wan L, Nagata T, Katahira M (2018) Influence of the DNA sequence/length and pH on deaminase activity, as well as the roles of the amino acid residues around the catalytic center of APOBEC3F. Phys Chem Chem Phys 20: 3109–3117.

71. Nakashima M, Ode H, Kawamura T, Kitamura S, Naganawa Y, et al. (2016) Structural Insights into HIV-1 Vif-APOBEC3F Interaction. J Virol 90: 1034–1047.

72. Hache G, Liddament MT, Harris RS (2005) The retroviral hypermutation specificity of APOBEC3F and APOBEC3G is governed by the C-terminal DNA cytosine deaminase domain. J Biol Chem 280: 10920–10924.

73. Chen Q, Xiao X, Wolfe A, Chen XS (2016) The in vitro Biochemical Characterization of an HIV-1 Restriction Factor APOBEC3F: Importance of Loop 7 on Both CD1 and CD2 for DNA Binding and Deamination. J Mol Biol 428: 2661–2670.

74. Schumann GG (2007) APOBEC3 proteins: major players in intracellular defence against LINE-1-mediated retrotransposition. Biochem Soc Trans 35: 637–642.

75. Schumann GG, Gogvadze EV, Osanai-Futahashi M, Kuroki A, Münk C, et al. (2010) Unique functions of repetitive transcriptomes. Int Rev Cell Mol Biol 285: 115–188.

76. Xie Y, Rosser JM, Thompson TL, Boeke JD, An W (2011) Characterization of L1 retrotransposition with high-throughput dual-luciferase assays. Nucleic Acids Res 39: e16.

77. Xiao X, Li SX, Yang H, Chen XS (2016) Crystal structures of APOBEC3G N-domain alone and its complex with DNA. Nat Commun 7: 12193.

78. Fang Y, Xiao X, Li SX, Wolfe A, Chen XS (2018) Molecular Interactions of a DNA Modifying Enzyme APOBEC3F Catalytic Domain with a Single-Stranded DNA. J Mol Biol 430: 87–101.

79. Marino D, Perkovic M, Hain A, Jaguva Vasudevan AA, Hofmann H, et al. (2016) APOBEC4 Enhances the Replication of HIV-1. PLoS One 11: e0155422.

80. Adolph MB, Ara A, Feng Y, Wittkopp CJ, Emerman M, et al. (2017) Cytidine deaminase efficiency of the lentiviral viral restriction factor APOBEC3C correlates with dimerization. Nucleic Acids Res 45: 3378–3394.

81. Solomon WC, Myint W, Hou S, Kanai T, Tripathi R, et al. (2019) Mechanism for APOBEC3G catalytic exclusion of RNA and non-substrate DNA. Nucleic Acids Res 47: 7676–7689.

82. Ziegler SJ, Hu Y, Devarkar SC, Xiong Y (2019) APOBEC3A loop 1 is a determinant for ssDNA binding and deamination. Biochemistry.

83. Bohn JA, DaSilva J, Kharytonchyk S, Mercedes M, Vosters J, et al. (2019) Flexibility in nucleic acid binding is central to APOBEC3H antiviral activity. J Virol.

84. Rathore A, Carpenter MA, Demir O, Ikeda T, Li M, et al. (2013) The local dinucleotide preference of APOBEC3G can be altered from 5’-CC to 5’-TC by a single amino acid substitution. J Mol Biol 425: 4442–4454.

85. Siu KK, Sultana A, Azimi FC, Lee JE (2013) Structural determinants of HIV-1 Vif susceptibility and DNA binding in APOBEC3F. Nat Commun 4: 2593.

86. Dang Y, Abudu A, Son S, Harjes E, Spearman P, et al. (2011) Identification of a single amino acid required for APOBEC3 antiretroviral cytidine deaminase activity. J Virol 85: 5691–5695.

87. Ohno S (1970) Evolution by gene duplication Springer-Verlag Heidelberg. Germany.

88. Athanassiou M, Hu Y, Jing L, Houle B, Zarbl H, et al. (1999) Stabilization and reactivation of the p53 tumor suppressor protein in nontumorigenic revertants of HeLa cervical cancer cells. Cell Growth Differ 10: 729–737.

89. Dull T, Zufferey R, Kelly M, Mandel RJ, Nguyen M, et al. (1998) A third-generation lentivirus vector with a conditional packaging system. J Virol 72: 8463–8471.

90. Bähr A, Singer A, Hain A, Vasudevan AA, Schilling M, et al. (2016) Interferon but not MxB inhibits foamy retroviruses. Virology 488: 51–60.

91. Russell RA, Wiegand HL, Moore MD, Schafer A, McClure MO, et al. (2005) Foamy virus Bet proteins function as novel inhibitors of the APOBEC3 family of innate antiretroviral defense factors. J Virol 79: 8724–8731.

92. Simm M, Shahabuddin M, Chao W, Allan JS, Volsky DJ (1995) Aberrant Gag protein composition of a human immunodeficiency virus type 1 vif mutant produced in primary lymphocytes. J Virol 69: 4582–4586.

93. Raiz J, Damert A, Chira S, Held U, Klawitter S, et al. (2012) The non-autonomous retrotransposon SVA is trans-mobilized by the human LINE-1 protein machinery. Nucleic Acids Res 40: 1666–1683.

94. Rose PP, Korber BT (2000) Detecting hypermutations in viral sequences with an emphasis on G --> A hypermutation. Bioinformatics 16: 400–401.

95. Jaguva Vasudevan AA, Perkovic M, Bulliard Y, Cichutek K, Trono D, et al. (2013) Prototype foamy virus Bet impairs the dimerization and cytosolic solubility of human APOBEC3G. J Virol 87: 9030–9040.

96. Robert X, Gouet P (2014) Deciphering key features in protein structures with the new ENDscript server. Nucleic Acids Res 42: W320–324.

97. Release S (2016) 2: Maestro, Schrödinger, LLC, New York, NY, 2017. Received: February 21: 2018.

98. Shi K, Carpenter MA, Banerjee S, Shaban NM, Kurahashi K, et al. (2017) Structural basis for targeted DNA cytosine deamination and mutagenesis by APOBEC3A and APOBEC3B. Nat Struct Mol Biol 24: 131–139.

99. Roshan U (2014) Multiple sequence alignment using Probcons and Probalign. Methods Mol Biol 1079: 147–153.

100. Waterhouse AM, Procter JB, Martin DM, Clamp M, Barton GJ (2009) Jalview Version 2--a multiple sequence alignment editor and analysis workbench. Bioinformatics 25: 1189–1191.

101. Bas DC, Rogers DM, Jensen JH (2008) Very fast prediction and rationalization of pKa values for protein-ligand complexes. Proteins 73: 765–783.

102. Jorgensen WL, Chandrasekhar J, Madura JD, Impey RW, Klein ML (1983) Comparison of simple potential functions for simulating liquid water. The Journal of chemical physics 79: 926–935.

103. D.A. Case VB, J.T. Berryman, R.M. Betz, Q. Cai, D.S. Cerutti, T.E. Cheatham, III, T.A. Darden, R.E. Duke, H. Gohlke, A.W. Goetz, S. Gusarov, N. Homeyer, P. Janowski, J. Kaus, I. Kolossváry, A. Kovalenko, T.S. Lee, S. LeGrand, T. Luchko, R. Luo, B. Madej, K.M. Merz, F. Paesani, D.R. Roe, A. Roitberg, C. Sagui, R. Salomon-Ferrer, G. Seabra, C.L. Simmerling, W. Smith, J. Swails, R.C. Walker, J. Wang, R.M. Wolf, X. Wu and P.A. Kollman (2014) AMBER 14. University of California, San Francisco.

104. Maier JA, Martinez C, Kasavajhala K, Wickstrom L, Hauser KE, et al. (2015) ff14SB: Improving the Accuracy of Protein Side Chain and Backbone Parameters from ff99SB. J Chem Theory Comput 11: 3696–3713.

105. Li P, Roberts BP, Chakravorty DK, Merz KM, Jr. (2013) Rational Design of Particle Mesh Ewald Compatible Lennard-Jones Parameters for +2 Metal Cations in Explicit Solvent. J Chem Theory Comput 9: 2733–2748.

106. Darden T, York D, Pedersen L (1993) Particle mesh Ewald: An N·log (N) method for Ewald sums in large systems. The Journal of chemical physics 98: 10089–10092.

107. Ryckaert J-P, Ciccotti G, Berendsen HJ (1977) Numerical integration of the cartesian equations of motion of a system with constraints: molecular dynamics of n-alkanes. Journal of computational physics 23: 327–341.

108. Hopkins CW, Le Grand S, Walker RC, Roitberg AE (2015) Long-Time-Step Molecular Dynamics through Hydrogen Mass Repartitioning. J Chem Theory Comput 11: 1864–1874.

109. Iwatani Y, Takeuchi H, Strebel K, Levin JG (2006) Biochemical activities of highly purified, catalytically active human APOBEC3G: correlation with antiviral effect. J Virol 80: 5992–6002.

110. Katoh K, Standley DM (2013) MAFFT multiple sequence alignment software version 7: improvements in performance and usability. Mol Biol Evol 30: 772–780.

111. Stamatakis A (2014) RAxML version 8: a tool for phylogenetic analysis and post-analysis of large phylogenies. Bioinformatics 30: 1312–1313.

112. Huson DH, Scornavacca C (2012) Dendroscope 3: an interactive tool for rooted phylogenetic trees and networks. Syst Biol 61: 1061–1067.

